# Ligand Binding Effects on Activation of Type-2 Angiotensin II Receptor

**DOI:** 10.1101/2025.01.22.634339

**Authors:** Ekrem Yasar, Segun Dogru, Erol Eroglu, Nazmi Yaras

## Abstract

1

The Angiotensin II type-2 receptor (AGTR2) is a G-protein coupled receptor (GPCR) in the Renin-Angiotensin System (RAS), mediating essential roles in cardiovascular regulation. Overactivation of AGTR2 causes to cardiovascular pathologies, ranging from hypertension to cancer. Despite its physiological importance, the molecular mechanisms driving AGTR2 activation remain poorly understood. Using long-timescale molecular dynamics (MD) simulations, we investigated the ligand binding effects of Angiotensin II (Ang II) and Angiotensin 1-7 (Ang 1-7) on AGTR2 in inactive and active states. Our results demonstrate that Ang II stabilizes the active-state of AGTR2 by reducing fluctuations in TM6 and Helix8, thereby promoting receptor activation. In contrast, Ang 1-7 modulates the inactive-state by enhancing flexibility in intracellular loops and Helix8, suggesting may a preparatory role for receptor activation. Key interactions, including hydrogen bonds (e.g., PRO7-THR125^3.33^ for Ang 1-7 and PHE8-LYS215^5.42^ for Ang II), salt bridges (e.g., ARG2-ASP279^6.58^ and ARG2-ASP297^7.32^), and hydrophobic contacts (e.g., TRP100^2.60^, MET128^3.36^), were identified as critical for receptor stability and ligand-specific dynamics. Analyses of conserved micro-switch motifs (CWxP, PIF, E/DRY, NPxxY) and intracellular distances revealed additional details about ligand-specific pathways contributing to AGTR2 activation and signal propagation. In addition, Dynamic Residue Interaction Network (DRIN) findings, aligned with micro-switch motif analyses, revealed that ligand binding impacts conserved regions, advancing our knowledge of AGTR2 activation mechanisms. These findings highlight the distinct yet complementary roles of Ang II and Ang 1-7 in regulating AGTR2 dynamics, offering new insights into GPCR activation and therapeutic opportunities for cardiovascular diseases.

**STATEMENT OF SIGNIFICANCE:** Understanding the intricate molecular mechanisms underlying the activation of G-protein coupled receptors (GPCRs), such as the Angiotensin II type-2 receptor (AGTR2), is vital for advancing therapeutic approaches targeting cardiovascular diseases. This study reveals that both Ang II and Ang 1-7 ligands stably bind to inactive and active-state AGTR2 structures throughout 1 microsecond (μs) long-timescale molecular dynamics (MD) simulations, influencing receptor dynamics and activation mechanisms within the protective arm of the Renin-Angiotensin System (RAS). While both ligands contribute to the stabilization of active-state conformations, Ang II demonstrates a more pronounced effect, potentially prolonging the receptor’s active state and enhancing intracellular signaling. These findings provide critical insights into the functional role of AGTR2 in RAS regulation and highlight its potential as a therapeutic target for conditions such as hypertension and other cardiovascular pathologies.

## 3 INTRODUCTION

The Renin-Angiotensin System (RAS) is a vital hormone system that its primary function lies in the maintenance of blood pressure and the regulation of fluid equilibrium (1–3). Beside vital roles, overactivation of the RAS may lead to adverse outcomes such as hypertension and kidney diseases (4–6). Consequently, pharmacological interventions targeting the RAS are employed to manage these disorders.

Angiotensin II (Ang II), synthesized by angiotensin-converting enzyme (ACE) through cleavage of the last two amino acids of inactive Angiotensin I (Ang I), consists of a polypeptide structure comprising eight amino acids and exists in its active form. The physiological effects of Ang II, such as aldosterone secretion, blood pressure regulation, water-electrolyte balance, vasoconstriction, catecholamine release, sodium, and water absorption, cell growth, and cardiac hypertrophy, are mediated by receptors within the RAS (7,8).

Various bioactive angiotensin peptides, including Angiotensin III (Ang III or Ang 2-8), Angiotensin IV (Ang IV or Ang 3-8), Alamandine, and Angiotensin 1-7 (Ang 1-7), are generated in the RAS by angiotensin-converting enzyme 2 (ACE2) and/or aminopeptidases through cleavage of certain amino acids from Ang II (9). Among these peptides, Ang 1-7 is produced through cleavage of phenylalanine (Phe), the terminal amino acid of Ang II. Ang 1-7, unlike Ang II, exerts blood vessel dilation, thereby reducing blood pressure, and is also believed to possess anti-inflammatory and anti-fibrotic properties (10–13). The RAS system exert its action through the receptors those are the Angiotensin II type 1 receptor (AGTR1), Angiotensin II type 2 receptor (AGTR2), proto-oncogene MAS receptor (MAS1), and MrgD receptor (14,15). Among these receptors, AGTR2 typically exhibits vasodilator effects, contributing to blood vessel dilation, and is implicated in anti-inflammatory and anti-proliferative processes. AGTR2 also plays a role in developmental processes and tissue repair (16–18).

Investigations into the properties and action mechanisms of RAS components have yielded significant insights into Ang peptides, related G protein-coupled receptors (GPCRs) and receptor blockers. Pharmacological inhibition of Ang II production and action represents a primary target for drug development due to its critical involvement in hypertension and other cardiovascular diseases (19). Recent evidence suggests that the beneficial effects of Ang 1-7 in the heart are mediated by the MAS1 (20). However, the signaling pathways underlying these effects in cardiomyocytes remain to be fully elucidated.

The three-dimensional (3D) structures of Ang II (PDB code: 1N9V) and Ang 1-7 (PDB code: 2JP8) peptides were determined using nuclear magnetic resonance (NMR) spectroscopy (21,22). The structural models of Angiotensin peptides and the receptors ensure the numerous studies have focused on modeling AGTR1 and AGTR2 (23–25). Utilizing the models proposed in structural modeling studies, researchers aim to understand how Angiotensin peptides bind to each receptor, the nature of their binding, and how binding alters the structure in active intracellular pathways. Integrating known functional data of AGTR1, AGTR2, and MAS1 receptors with predictions from molecular docking of Ang peptides using structural protein modeling techniques, sequence alignments, and molecular dynamics (MD) simulations, it is anticipated that conserved binding roles for Ang peptides at these receptors can be identified and potential protein interactions elucidated (26–28).

R. A. Santos group used I-TASSER and homology modeling to generate 3D structures of AGTR1, AGTR2, and MAS1, followed by MD simulations to evaluate ligand binding stability through molecular docking with Angiotensin peptides (24,29). However, while these studies provided valuable insights, their short simulations offered limited evidence on ligand-induced receptor dynamics, differing from our approach by focusing on comprehensive receptor activation analyses through advanced and long-duration simulations.

The crystal structures of AGTR1 and AGTR2 in ligand-bound states (30–33) and the recent cryo-EM structures of MrgD and AGTR1 (34,35) provided valuable insights into the binding mechanisms and activation of RAS receptors. These studies revealed key interactions of Angiotensin peptides with RAS receptors, advancing our understanding of receptor dynamics and facilitating structure-based drug discovery for cardiovascular diseases. While the structural and functional roles of AGTR2, particularly in the protective arm of the RAS, have been highlighted, further research is needed to elucidate its activation mechanisms and dynamics.

In this study, MD simulations were employed to investigate the structural dynamics of AGTR2, a member of the class-A GPCR family, with a focus on its interactions with Angiotensin peptides. The primary goal was to gain novel insights into the receptor’s activation mechanism and validate the general GPCR activation pathway driven by ligand binding in the context of AGTR2. Our findings contribute to a deeper understanding of the molecular mechanisms underlying the protective arm of the RAS, shedding light on its functional dynamics.

## 4 MATERIALS AND METHODS

### 4.1. Computational Design

The structures of Ang II (PDB code: 1N9V) (21) and Ang 1-7 (PDB code: 2JP8) (22) ligands obtained using the NMR spectroscopy method were taken from the protein databank (PDB) and used in our study.

This active-like structure of AGTR2, which was determined by X-ray crystallography method and bound to Ang II peptide, was taken from PDB (36) (PDB code: 6JOD) (33). Since there was no 3D structure in the inactive state obtained by experimental methods, the inactive-like AGTR2 structure was taken from the GPCR database (GPCRdb) (37,38). These structures can be found in Fig S1 in the supplemental figures.

All missing residues in the structure files were prepared using the MODELLER web server (39). to prepare only the structure of the desired molecule for use. The protonation states of all titratable structures were determined at pH: 7.0 (neutral) using the H++ web server (40), and missing hydrogen atoms were completed. The structures obtained from the H++ web server was used for molecular docking.

### 4.2. Molecular Docking

By using blind docking as a molecular docking method, ligand binding poses with the highest binding affinity to the inactive-state and active-state structures of the receptor were determined. For molecular docking, MDockPeP: ab-initio protein-peptide docking program using a statistical potential-based scoring function (ITScorePeP) (41) was preferred. Ang II and Ang 1-7 ligands were docked separately to both inactive-state and active-state structures of AGTR2 and initial poses were obtained for simulations. Since the active structure of AGTR2 was obtained from the PDB database (PDB code: 6JOD) and this crystal structure contains the Ang II bound receptor structure (33), this structure was preferred in simulations for Ang II bound active-state AGTR2. Besides, Ang II was removed from this structure and docking was also performed to determine docking score. After all protein-peptide docking process, the poses with the lowest ITScorePeP score were selected for MD simulations.

### 4.3. Molecular Dynamics Simulation

A total of 6 different structures were used for Molecular Dynamics (MD) simulations, including AGTR2-Ang II (inactive-state and active-state), AGTR2-Ang (1–7) (inactive-state and active-state), protein-ligand complex structures, and AGTR2 in inactive and active states without ligand binding (Apo-AGTR2).

All protein structures used were placed on the membrane consisting of a lipid bilayer. The orientation of the constructs concerning the membrane was determined using the Positioning of Proteins in Membrane (PPM) server of the Orientations of Proteins and Membranes (OPM) database (42). All structures were assembled using the Membrane Builder package of the CHARMM-GUI program (43–45) into a lipid bilayer cocktail consisting of 1-palmitoyl-2-oleoyl-sn-glycero-3-phosphocholine (POPC), 1-palmitoyl-2-oleoyl-snglycero-3-phosphoethanolamine (POPE) and cholesterol (CHL) using POPC: POPE: CHL=2:2:1 ratio. For MD simulations, all of the ligand-protein-membrane complex systems formed with the membrane-inserted structures were placed in 90×90×90 nanometer (nm) simulation boxes, added TIP3P type water molecules (46) and neutralized with 0.15 mM (millimolar) NaCl. In addition to receptor and ligands, each system contains approximately 253 lipids, ∼63,000 water molecules, 50-60 sodium (Na^+^) and 60-70 chlorine (Cl^-^) ions, and approximately 100,000 atoms.

Charmm36m force field (47) was used for the complex structures and all systems were subjected to MD simulations using computer systems with NVIDIA RTX A4000 GPU graphics cards with GROMACS 2020.5 version software package (48).

Before MD simulations, each system setup was relaxed to eliminate possible steric collapses by performing 5000 steps of conjugate gradient minimization. Harmonic constraining force constants of 10 kcal/mol/Å2 were applied to receptor and ligand residues as well as lipid molecules. In the equilibration step, the harmonic constraint force constant for each system was set to 5 kcal/mol/Å2. In all systems, the temperature was gradually raised from 0 K to 310 K and equilibrated over 10 ns by applying a Nose-Hoover thermostat (49) with a temperature coupling constant of 1.0 ps in the NVT section, followed by a quasi-isotropic Berendsen barostat (50) using a constant pressure group (NPT), and the pressure was kept constant throughout the MD simulation runs using a quasi-isotropic Parrinello-Rahman barostat (51). Simulation runs of 2 x 1000 ns duration for each system were performed using periodic boundary conditions and a time step of 2 fs. A separate replicate was made for each simulation (Fig S11), and one of them was randomly selected for each analysis of both bound and unbound ligand structures. During the MD simulations, the Particle Mesh Ewald (PME) method (52) was applied to calculate the exact electrostatic energy of a unit cell in a macroscopic lattice of repeating images. All bonds were analyzed using the SHAKE algorithm (53) and were constrained accordingly, ensuring the stability and accuracy of the simulations.

### 4.4. Data Analysis

Trajectory analyses, including root-mean-square deviation (RMSD), root-mean-square fluctuation (RMSF), and center of mass (COM) distance measurements, were performed using the analytical tools provided within the GROMACS gmx packages (54). To determine 2-dimensional (2D) interactions between the ligand and receptor, the Maestro program (Schrödinger Maestro, 2020) was utilized during a 1-month free demo provided by Schrödinger (55).

Additionally, ligand-protein interactions including hydrogen bonds, salt bridges, and hydrophobic interactions, were thoroughly analyzed. Throughout the simulations, hydrogen bond analyses (2000 frames) and salt bridge analyses (10000 frames) were conducted using the VMD 1.9.3 program (56). In addition, ligand-protein interactions were also visualized through movies using VMD program. Hydrophobic interaction occupancy (%) was assessed for 2000 frames using Python tools including the MDAnalysis program package (57), with the corresponding scripts available on GitHub (https://github.com/eygpcr/hydrophobic_interactions).

In addition to trajectory analysis tools, several advanced methods have been used to analyse molecular dynamics (MD) simulations to assess the structural and dynamic changes induced by ligand binding in AGTR2. These included the MDM-TASK web server (58,59), which offers techniques such as Communication Propensity (CP), Dynamic Cross-Correlation (DCC), Betweenness Centrality (BC), Dynamic Residue Interaction Network (DRIN), and Principal Component Analysis (PCA). A theoretical overview of these methods can be found in the supplementary material of our previous publication (60).

Considering the common activation mechanism proposed for class-A GPCRs, the gmx_distance tool was used to analyze distance changes between transmembrane helices in the intracellular region, known as the hydrophobic lock. Additionally, phi (ϕ), psi (ψ) and chi (χ) dihedral angle changes in the PIF, E/DRY, CWxP, and NPxxY micro-switch regions of AGTR2, based on ligand binding, were examined using Python applications including the MDAnalysis program package have been added to Github (https://github.com/eygpcr/microswitches_dihedral_angles).

The trajectory files for all AGTR2 simulations have been uploaded to the GPCRmd database (61) under PDB code: 6JOD and Uniprot ID: P50052, and all data is publicly accessible through the GPCRmd platform (https://www.gpcrmd.org/dynadb/search/).

## 5 RESULTS

### 5.1. Ligand-Receptor Interactions

Molecular docking results revealed that the docking scores of the Ang 1-7 ligand to AGTR2 were similar for both inactive-state (−161.0) and active-state (−179.6) structures, while Ang II showed a better docking score for the inactive-state structure (−163.4) and the best score for the active-state structure (−208.1). The 2D ligand-protein binding interactions corresponding to these docking poses are shown in Fig S2 and S3. During the 1000 ns MD simulations, both Ang II and Ang 1-7 ligands remained stably bound to inactive-state and active-state AGTR2 structures (Supplementary Movies S1-S4). Distance analyses showed that the distance between the last amino acid of Ang II (PHE8) and the amino acid MET128^3.36^ (62) of AGTR2, which is thought to play a key role in the activation of AGTR2 by hydrophobically binding (33), remained stable in both inactive-state and active-state structures, while the distance between the last amino acid of Ang 1-7 (PRO7) and MET128^3.36^ was stable in the inactive structure but increased by 4 Å in the active-state structure compared to the initial docking pose, as illustrated in Fig S5.

#### 5.1.1. Hydrogen bonding

The hydrogen bond analysis revealed distinct binding patterns of Ang 1-7 and Ang II to inactive-state and active-state AGTR2 structures as illustrated in Fig 1a. For Ang 1-7, the number of hydrogen bonds contributing to binding was higher in the inactive-state structure than in the active-state structure, and these bonds were maintained throughout the simulation. Key hydrogen bonds in the inactive-state structure included PRO7-THR125^3.33^, ARG2-ASP297^7.32^, ARG2-ASP279^6.58^, and PRO7-LYS215^5.42^, while the primary contributor in the active-state structure was the ASP1-GLU188 bond. Notably, no common hydrogen bonds were observed between Ang 1-7 binding to inactive-state and active-state structures.

**FIGURE 1.**
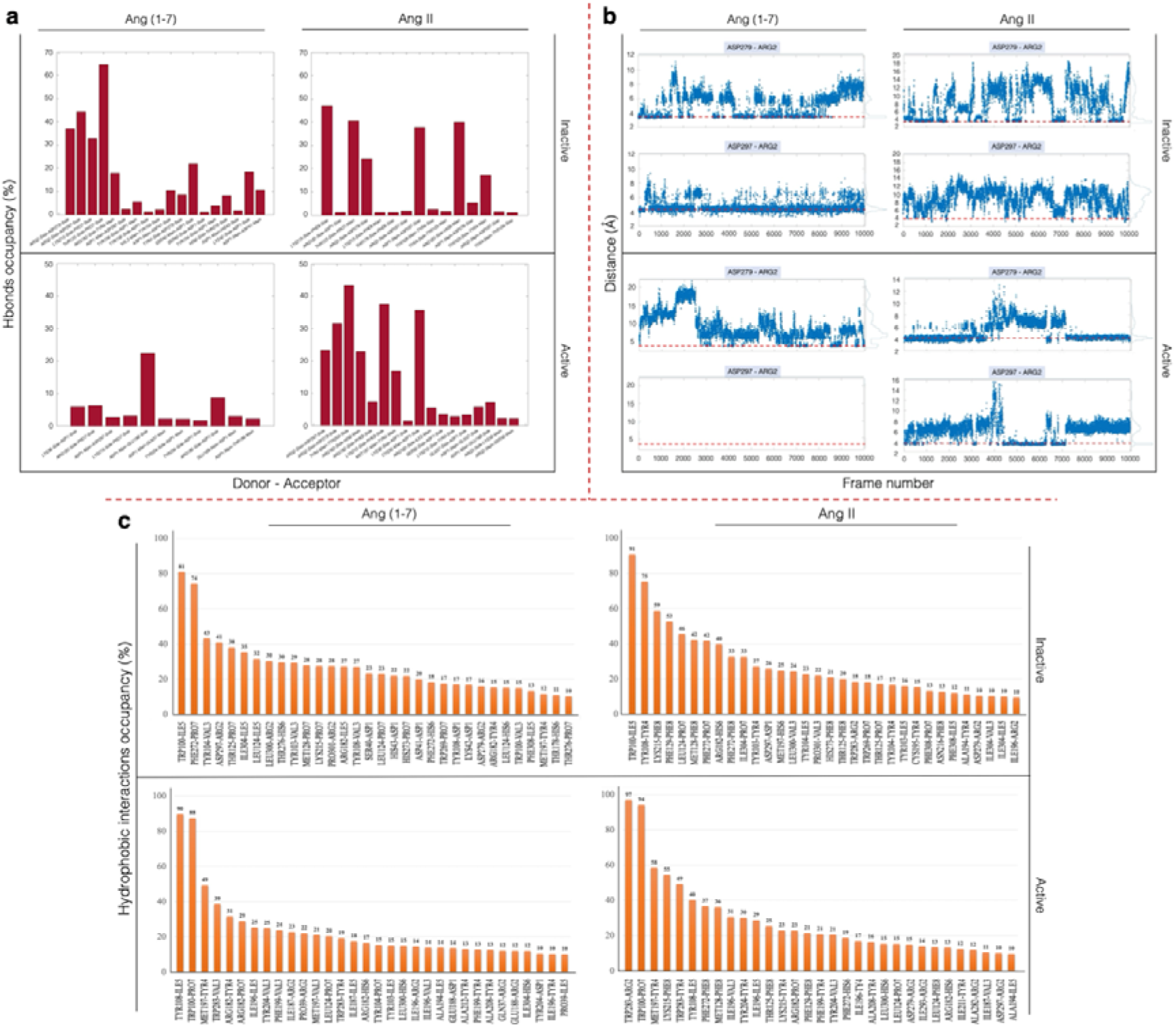
**a)** Hydrogen-bond occupancies (%), b) Distance fluctuations (Å) of key salt bridges (ARG2-ASP279^6.58^ and ARG2-ASP297^7.32^) and c) Hydrophobic interaction occupancies (%) between the ligands (Ang 1-7 and Ang II) and AGTR2 in inactive-state and active-state structures during the 1000 ns simulation.

In contrast, Ang II exhibited more hydrogen bonds in the active-state structure compared to the inactive-state structure. Key bonds in the inactive-state structure included PHE8-LYS215^5.42^, PRO7-THR125^3.33^, HIS6-ARG182^4.64^, ASP1-ASP297^7.32^, and ARG2-ASP279^6.58^ residue pairs, whereas the active-state structure showed contributions from TYR4-TYR204^5.31^, PHE8-LYS215^5.42^, ASP1-CYS35, ARG2-ASP279^6.58^, ARG2-ASP297^7.32^, and HIS6-ARG182^4.64^. Common hydrogen bonds for Ang II binding to both structures included PHE8-LYS215^5.42^, ARG2-ASP279^6.58^, and HIS6-ARG182^4.64^. Additionally, PRO7-THR125^3.33^ and ARG2-ASP279^6.58^ were shared bonds in Ang 1-7 and Ang II binding to the inactive-state structure. Furthermore, in the active-state structure, Ang 1-7 formed a hydrogen bond via PRO7-LYS215^5.42^, while Ang II utilized PHE8-LYS215^5.42^ for the same interaction. All hydrogen bonds formed during the interactions of the AGTR2 in its inactive and active states with the ligands Ang 1-7 and Ang II throughout the simulation are summarized in Table S1 and S2.

#### 5.1.2. Salt bridge

Salt bridge analysis revealed that the key residues contributing to the binding of both Ang 1-7 and Ang II to inactive-state and active-state AGTR2 structures were ARG2-ASP279^6.58^ and ARG2-ASP297^7.32^ residue pairs, as shown in Fig 1b. In the inactive-state structure, Ang 1-7 primarily relied on the ARG2-ASP297^7.32^ interaction, which was consistently maintained throughout the simulation, while in the active-state structure, this salt bridge was absent, and interaction depended on ARG2-ASP279^6.58^.

For Ang II, binding to the inactive-state structure was primarily mediated by ARG2-ASP279^6.58^, although neither salt bridge was maintained throughout the simulation. In the active-state structure, the ARG2-ASP279^6.58^ bond initially contributed the most, but between 400–700 ns, the bond was temporarily broken, and ARG2 formed a salt bridge with ASP297^7.32^ instead. Thus, the salt bridge between ligand and protein was maintained throughout the simulation.

#### 5.1.3. Hydrophobic interaction

The hydrophobic interactions contributing to ligand binding and stability in AGTR2 structures during simulations, as illustrated in Fig 1c, revealed both common and distinct patterns for Ang II and Ang 1-7 ligands in inactive and active states. Key receptor residues TRP100^2.60^, TYR103^2.63^, TYR104^2.64^, LEU124^3.32^, MET128^3.36^, ILE196^45.51^, MET197^45.52^, and PHE272^6.51^ were found to engage in hydrophobic interactions with both ligands across all binding states (Fig S2 and S3).

Specific hydrophobic interactions unique to each ligand and binding state were also identified. For Ang 1-7 ligand, ILE5-TRP100^2.60^ and PRO7-PHE272^6.51^ residue pairs were critical in the inactive structure, while ILE5-TYR108 and PRO7-TRP100^2.60^ contributed in the active structure. For Ang II ligand, ILE5-TRP100^2.60^ and TYR4-TYR108 residue pairs were important in the inactive structure, whereas ARG2-TRP283^6.62^ and PRO7-TRP100^2.60^ dominated in the active structure. Among these, the TRP100^2.60^ residue was a common contributor to the binding of both ligands in all states, highlighting its pivotal role in ligand-receptor interactions. These hydrophobic forces were consistently maintained throughout the MD simulations, underscoring their importance in stabilizing ligand binding.

### 5.2. Structural Dynamic Changes

The conformational and structural dynamic changes in AGTR2 affected by Ang II and Ang 1-7 ligands during MD simulations were analyzed, with results presented in Fig 2. RMSD and RMSF graphs illustrate the structural stability and flexibility of inactive-state and active-state AGTR2 with and without ligand binding.

**FIGURE 2.**
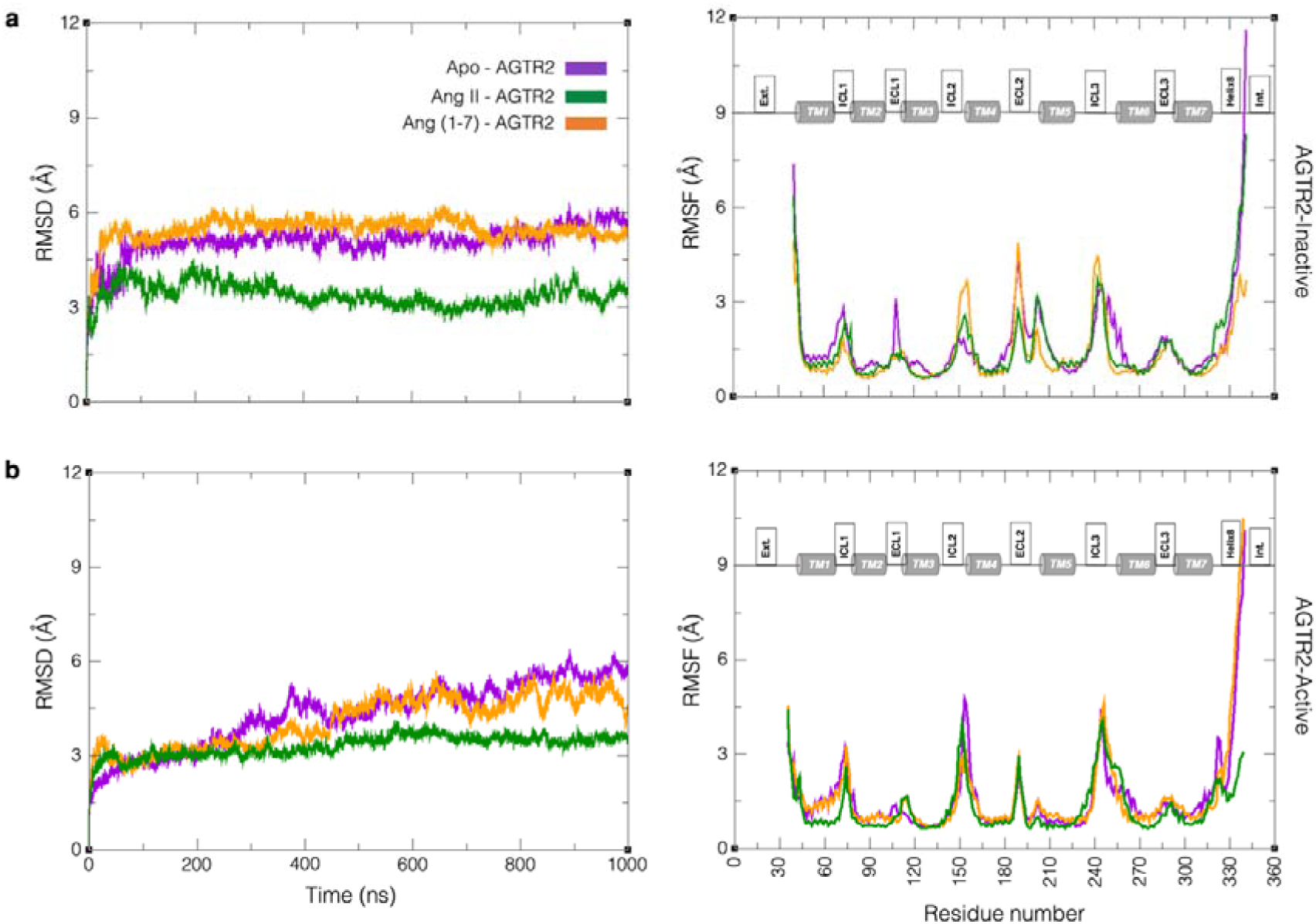
**a)** RMSD and RMSF plots of the inactive-state and b) the active-state AGTR2 structures with Apo (purple), Ang II-bound (green), and Ang (1–7)-bound (orange) states over the 1000 ns MD simulation.

For the inactive-state, the RMSD analysis showed that Apo-AGTR2 and Ang (1–7)-bound AGTR2 exhibited similar stability (∼5.5-6 Å), while the Ang II-bound AGTR2 structure displayed a lower RMSD (∼3 Å), indicating a more stable receptor structure compared to the ligand-free state. RMSF analysis revealed that Ang 1-7 binding caused increased fluctuations in ICL2, ECL2, and ICL3 compared to Apo-AGTR2, while Ang II binding caused fluctuations in ICL2, lower than those observed with Ang 1-7. Both ligands reduced fluctuations in ECL1, the ligand-binding site, compared to Apo-AGTR2. In the active-like state, RMSD plots showed similar stability for the Ang (1–7)-bound and Apo-AGTR2 structures, with an initial RMSD of ∼3 Å increasing to ∼5.5-6 Å after 500 ns. The Ang II-bound AGTR2 structure maintained a consistent RMSD of ∼3 Å, indicating greater structural stability compared to the ligand-free state. RMSF analysis revealed that both Ang 1-7 and Ang II binding reduced fluctuations in ICL2 compared to Apo-AGTR2, while other regions remained largely unaffected.

The RMSD changes of the transmembrane (TM) helices and Helix8 in AGTR2, with and without ligand binding, are presented in Fig S4. For the inactive-state AGTR2 structure, Ang II binding led to increased stability in TM1, TM3, TM4, TM6, and TM7 compared to the Apo-AGTR2 state, with the most pronounced stabilization observed in TM1 and TM6. No significant changes were observed in TM2 and TM5. In the active-state structure, Ang II binding stabilized TM1, TM6, and Helix8, with Helix8 showing the highest stability. However, the stabilization of TM6 persisted only for the first 400 ns of the simulation.

With Ang 1-7 binding to the inactive-state AGTR2, TM3 and TM4 became more stable compared to Apo-AGTR2, while Helix8 exhibited decreased stability. No changes were observed in TM1, TM2, TM5, TM6, or TM7. For the active-state AGTR2 structure, Ang 1-7 binding caused no significant changes in the stability of any TM helices or Helix8 compared to the Apo-AGTR2 state.

#### 5.2.1. Communication Propensity (CP)

CP of a residue pair depends on the fluctuations in their inter-residue distance. Smaller, low-intensity fluctuations lead to faster communication, whereas larger fluctuations result in slower communication (63). The results of CP analyses for residue pair interactions in AGTR2 with and without ligand binding, in both inactive and active states, are shown in Fig 3a. In the inactive-state, Apo-AGTR2 displayed high communication tendencies in ECL1, ECL2, ICL1, and ICL3. Upon binding of Ang 1-7, communication tendencies decreased in ECL1, ECL2, and ICL1, while increasing in ICL2 and ICL3. With Ang II binding, tendencies decreased in ECL1, ECL2, and ICL3, increased in ICL2, and remained unchanged in ICL1 compared to Apo-AGTR2.

**FIGURE 3.**
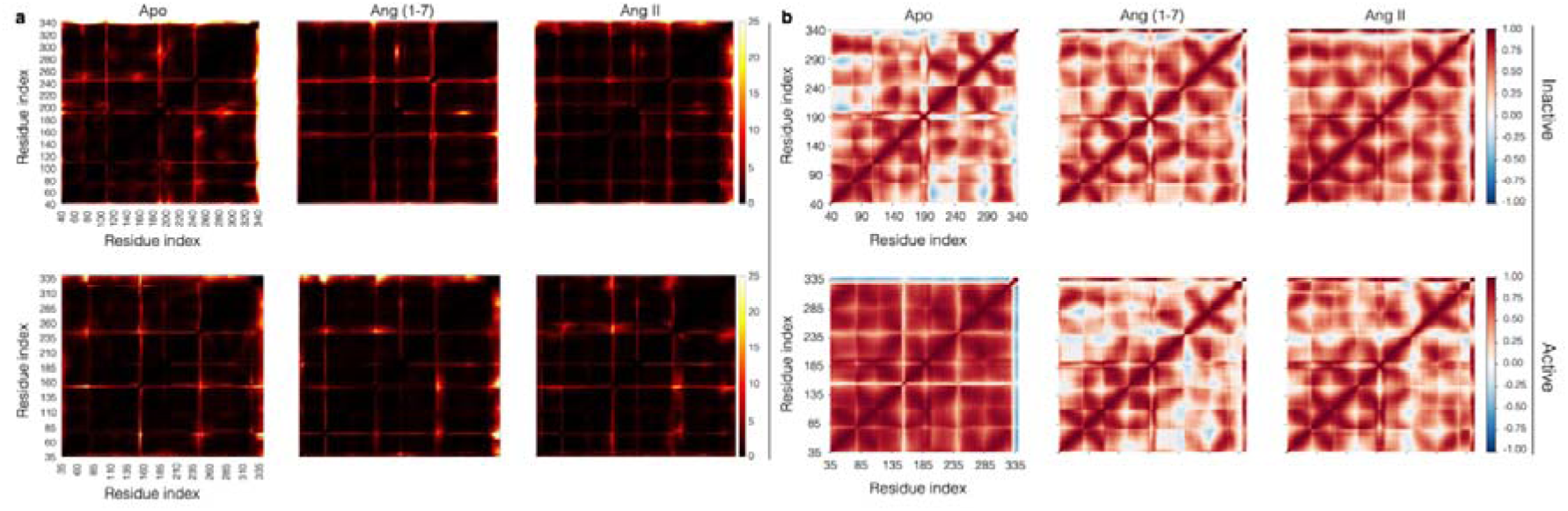
**a)** Communication propensity (CP) maps and b) Dynamic cross-correlation (DCC) maps of Apo-AGTR2, Ang (1–7)-AGTR2, and Ang II-AGTR2 complex structures both inactive and active states.

In the active-state, Apo-AGTR2 exhibited high communication tendencies in ECL2, ICL1, ICL2, ICL3, and Helix8. When Ang 1-7 was bound, tendencies decreased in ICL2, ICL3, and Helix8, increased in ICL1, and remained unchanged in ECL2. With Ang II binding, communication tendencies decreased in Helix8, increased in ICL1, and remained unchanged in ECL2, ICL2, and ICL3 compared to Apo-AGTR2.

#### 5.2.2. Dynamic Cross Correlation (DCC)

DCC analysis was conducted on the trajectory files from MD simulations of AGTR2 in its Ang (1–7)-bound, Ang II-bound, and ligand-free forms. The resulting DCC maps may provide insights into interactions between different regions of the receptor throughout the simulations. The results of DCC analysis for residue pair interactions in AGTR2 before and after ligand binding, in both inactive-like and active-like states, are shown in Fig 3b.

When the mutual and internal correlations of the TM helices of the receptor and the components, for the inactive-state Apo-AGTR2, mutual correlations between TM1-TM5, TM1-ICL3, TM1-ECL3, TM2-TM5, TM2-ICL3, TM2-ECL3, TM5-TM7, and ECL3-TM7 increased upon Ang 1-7 binding and further increased with Ang II binding. Correlations between TM3-ECL2, TM5-ECL2, ICL3-ECL2, and ECL3-ECL2 decreased with Ang 1-7 binding but increased with Ang II binding. Additionally, correlations of TM1, TM2, TM3, and TM6 with Helix8 decreased with Ang 1-7 but increased with Ang II binding.

In the active-state Apo-AGTR2, a very high correlation was observed among all components in the ligand-free state. Upon Ang 1-7 binding, decreases were noted in correlations between TM1-TM3, TM1-TM5, TM1-TM6, TM1-ICL3, TM1-ECL3, TM2-TM5, TM2-ICL3, TM3-ECL3, TM5-ECL3, TM4-ICL3, ECL2-ICL3, ECL3-TM7, and ICL3-TM6. For Ang II binding, reductions were seen in correlations between TM1-TM5, TM1-TM6, TM2-TM5, TM4-TM5, ECL2-TM5, TM5-TM6, TM5-Helix8, and TM6-Helix8.

#### 5.2.3. Principal Component Analysis (PCA)

PCA is a statistical method that diagonalizes the covariance matrix, removing linear correlations among atomic coordinates. The first few principal components, representing the largest fluctuations, can be analyzed through scatter plots to visualize how the system explores the configuration space. The PCA of AGTR2 fluctuations in inactive-state and active-state structures, in the Apo-state and with Ang 1-7 and Ang II ligands bound, is shown in Fig 4.

**FIGURE 5.**
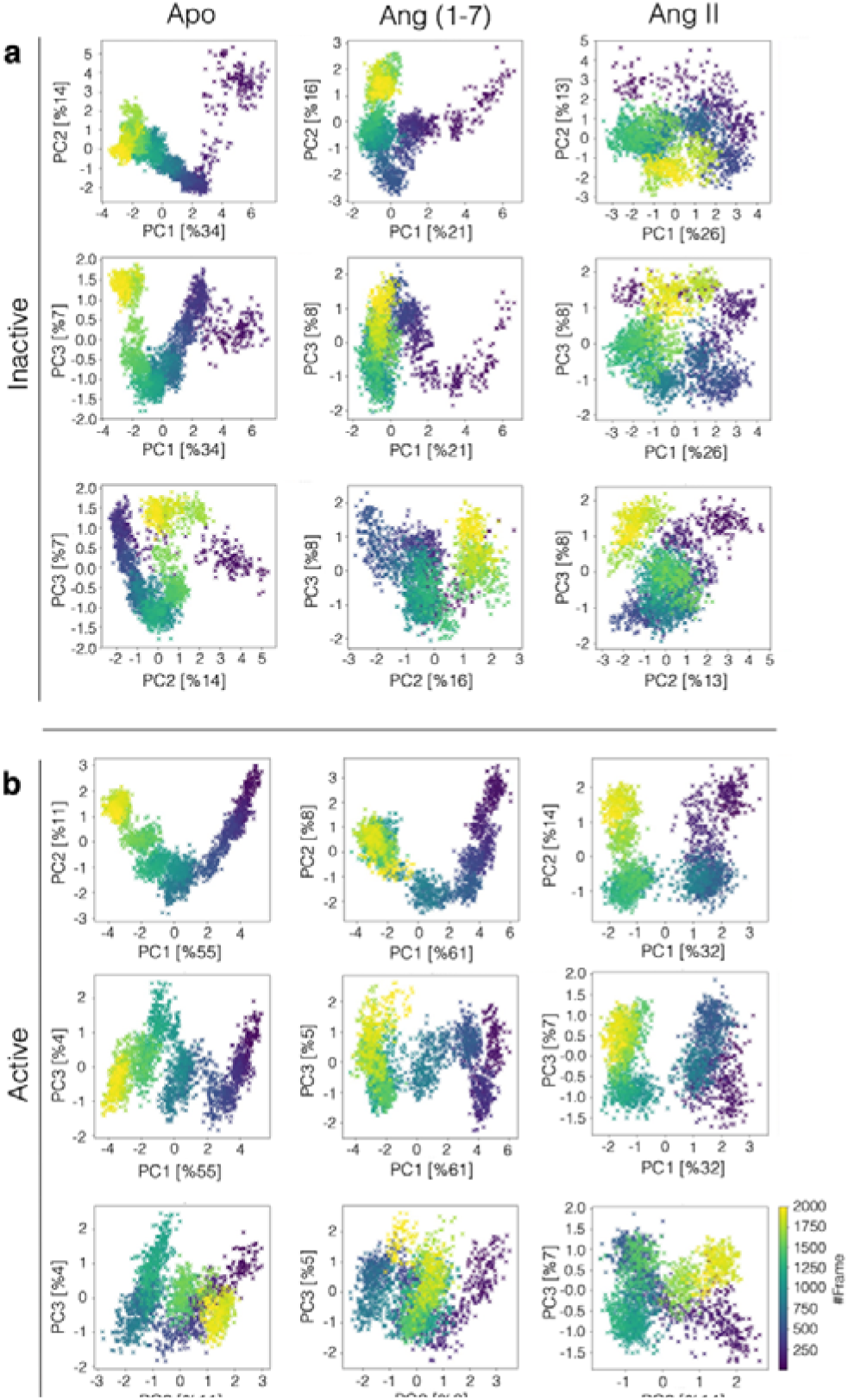
PCA analyses for AGTR2. PC1-PC2, PC1-PC3, PC2-PC3 exchange plots of Apo-AGTR2, Ang (1–7)-AGTR2 and AngII-AGTR2 complex structures in a) inactive-state and b) active-state.

For the inactive-state AGTR2 structure, the first principal component (PC1) explains 34% of the structural variations in the Apo-state but only 21% and 26% in the Ang 1-7- and Ang II-bound states, respectively, indicating that structural variations are not solely captured by PC1 upon ligand binding. The second principal component (PC2) accounts for 14% of the variations in the Apo-state, 16% in the Ang 1-7-bound state, and 13% in the Ang II-bound state. The third principal component (PC3) explains 7-8% of the variations across all states. The scatter plot of PC1 versus PC2 reveals three sampled states, with similar residue movements in the Apo-state and Ang 1-7-bound state, while the Ang II-bound state exhibits distinct movements.

For the active-state AGTR2 structure, PC1 contributes significantly to the structural variations, explaining 55% in the Apo-state, 61% in the Ang 1-7-bound state, and 32% in the Ang II-bound state. PC2 accounts for 11%, 8%, and 14% of the variations, respectively, while PC3 explains 4-7%. The PC1 versus PC2 scatter plot shows three distinct sampled states, with residue movements along eigenvectors 1 and 2 being similar for the Apo-state and Ang 1-7-bound state but differing significantly in the Ang II-bound state. These findings indicate ligand-specific effects on AGTR2 dynamics in both conformational states.

#### 5.2.4. Dynamic Residue Interaction Network (DRIN)

In graph theory, the shortest path represents the reachability between residues, defined as the minimum number of connections needed between residues i and j. Betweenness centrality (*BC*) quantifies how often a residue lies on these shortest paths within a residue interaction network. *BC* and shortest path length (*L*) were calculated to analyze residue connectivity during simulations of Apo-AGTR2, Ang II-AGTR2 and Ang (1–7)-AGTR2 complex systems. To evaluate the effects of Ang II and Ang (1–7) binding to AGTR2, the differences in average BC and *L* values between ligand-bound and Apo-AGTR2 states were determined.

The Δ*BC* and Δ*L* plots, illustrating the effects of Ang 1-7 and Ang II binding on AGTR2 receptor dynamics and signaling in inactive and active-states, are shown in Fig 5. Residues with a significant increase/decrease in mean *BC* and *L* values upon Ang 1-7 and Ang II binding are listed in Table S3 and S4, respectively. In addition, the *BC* increase changes in the inactive and active states upon ligand binding are visualized by the snake diagram in Fig S6. For the inactive-state AGTR2 structure, Ang 1-7 binding shows to notable changes in the *BC* and *L* values of residues in the intracellular loop regions and the TM6 helix. This binding enhances the signaling roles of ICL1, ICL2, ICL3, TM6, and Helix8 (+Δ*BC*), while Δ*L* values are negative for all components except ICL2. In contrast, Ang II binding affects all loop regions and TM helices, significantly increasing their signaling roles (+Δ*BC*) while Δ*L* values are consistently negative.

**FIGURE 5.**
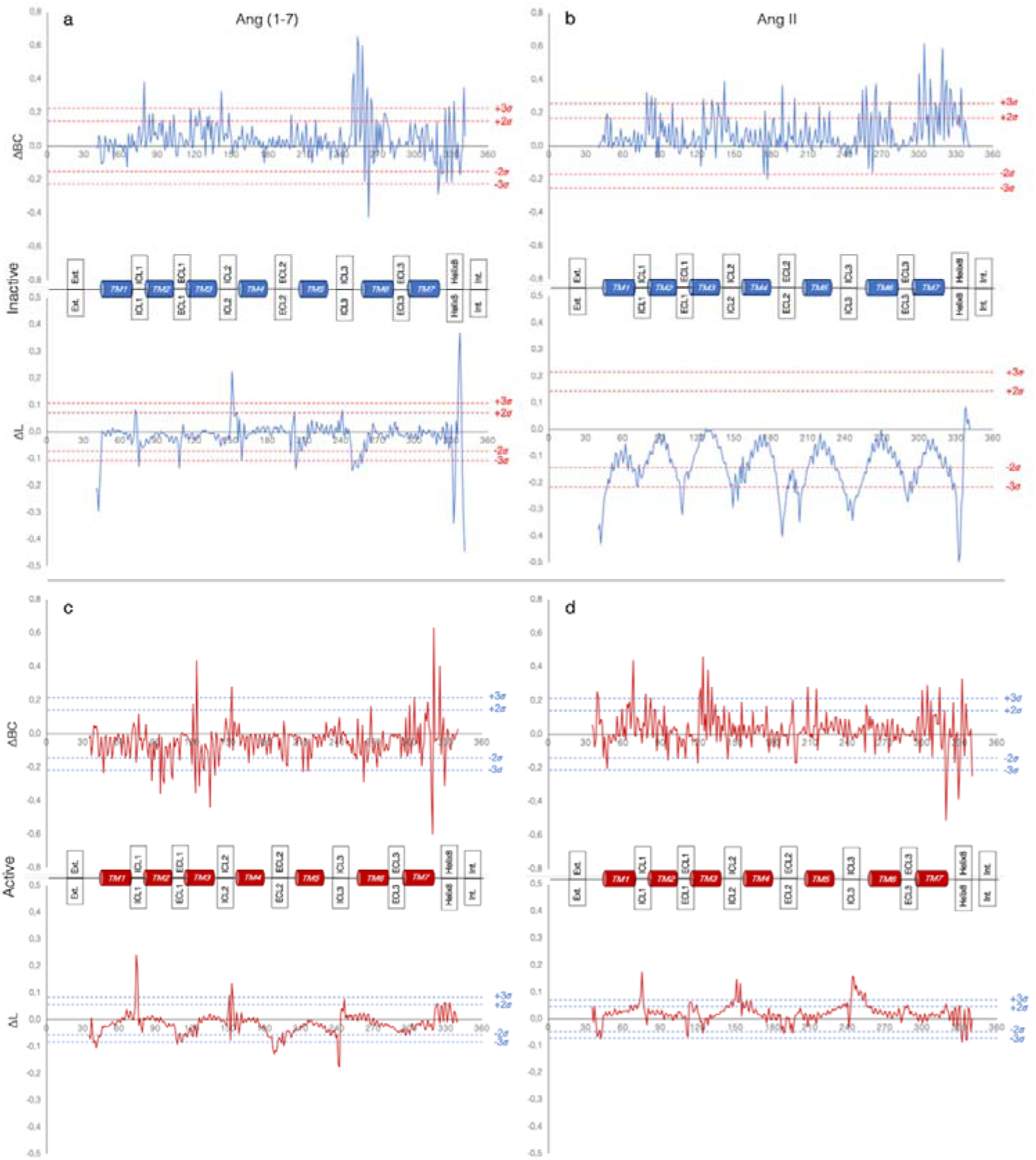
DRIN analyses for AGTR2 in the inactive-state. Plots of Δ*BC* and Δ*L* change upon binding of a) Ang 1-7 and b) Ang II ligands to inactive-state AGTR2. DRIN analyses for AGTR2 in the active-state. Plots of Δ*BC* and Δ*L* change upon binding of c) Ang 1-7 and d) Ang II ligands to active-state AGTR2.

For the active-state AGTR2 structure, Ang 1-7 binding leads to notable changes in the *BC* and *L* values of TM2, TM3, TM7, and Helix8. It enhances the signaling roles of TM3, ICL2, and Helix8 (+Δ*BC*) while reducing signaling in TM2, TM3, and TM7 (−Δ*BC*). Δ*L* values are positive for ICL1 and ICL3 but negative for other loops. Ang II binding, however, causes the largest changes in TM1, TM3, TM5, TM7, and Helix8. It enhances the signaling roles of TM1, ICL1, TM3, TM5, and TM7 (+ΔBC) while reducing the signaling role of Helix8 (-Δ*BC*). Additionally, Δ*L* values for ICL1, ICL2, and ICL3 are positive.

Comparison of Ang 1-7 binding in inactive and active states shows that positive *BC* values, contributing to signaling, are higher in the inactive state, while negative *BC* values are higher in the active state. Similarly, for Ang II binding, positive *BC* values enhance signaling in both states, but negative Δ*L* values are more pronounced in the inactive state, highlighting its greater role in stabilizing receptor dynamics.

### 5.3. Common Activation Mechanism

The generally accepted activation mechanism for class-A GPCRs involves ligand-induced changes in intracellular distances between key receptor components. This process primarily affects the helices forming the hydrophobic lock between TM3 and TM6, originating from the ligand-binding site. Additionally, conserved micro-switch residues play a crucial role in signaling by undergoing angular changes that contribute to receptor activation.

#### 5.3.1. Hydrophobic-Lock

The alterations in intracellular distances between TM helices (TM3, TM5, TM6, and TM7) of AGTR2 in inactive-state and active-state structures, with and without ligand binding, are shown in Fig 6.

**FIGURE 6.**
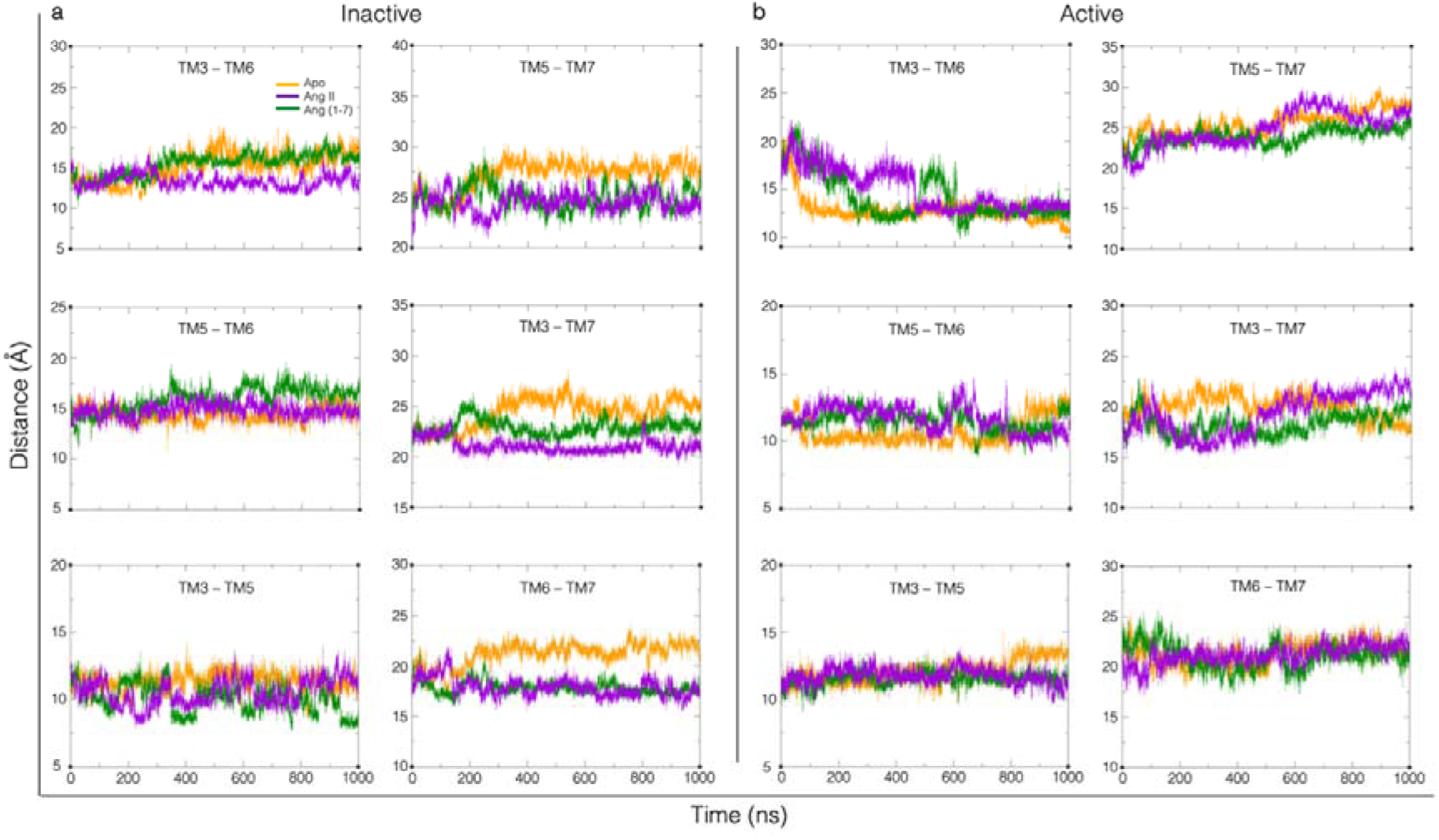
Hydrophobic-lock distance analyses for AGTR2. a) In the inactive-state and b) the active-state, variations in intracellular distances between TM3-TM6, TM5-TM7, TM5-TM6, TM3-TM7, TM3-TM5, and TM6-TM7 of Apo-AGTR2, Ang (1–7)-AGTR2, and Ang II-AGTR2 complex structures are depicted through graphical representations of changes over MD simulations.

For the inactive-state AGTR2, with the binding of Ang 1-7 showed in an increase in the TM3-TM6 distance after 250 ns, rising from ∼15 Å to ∼18 Å, and then exhibited similar change to that observed in the Apo-state. In contrast, Ang II binding did not alter the TM3-TM6 distance, which remained constant at around ∼15 Å throughout the simulation. Other TM helix pairs showed varying behaviors. For the active-state AGTR2, in the Apo-state observed a ∼7.5 Å decrease from around 20 Å in TM3-TM6 distance after 60 ns, which further decreased by 2.5 Å after 800 ns. With Ang 1-7 binding, the TM3-TM6 distance decreased after 300 ns, temporarily increased by 3-4 Å between 500-600 ns, and then returned to ∼7.5 Å. With also Ang II binding, the TM3-TM6 distance remained stable at around ∼17.5 Å during the first 500 ns before decreasing by ∼5-6 Å to ∼12.5 Å, where it remained stable until the end of the simulation.

#### 5.3.2. Micro-Switch Motifs and Dihedral Angles

Changes in the phi (ϕ), psi (ψ), and chi (χ) dihedral angles of residues in conserved micro-switch motifs (CWxP, E/DRY, PIF, NPxxY) were analyzed to determine structural alterations mediating AGTR2 activation upon Ang 1-7 and Ang II binding. The results, highlighting dynamic changes contributing to receptor activation, are depicted in Fig S7-S10.

##### PIF Motif

No significant angular changes were observed for PRO223^5.50^ and ILE132^3.40^ in any state. For PHE265^6.44^, the psi angle showed a notable increase of ∼10° with Ang II binding and ∼30° with Ang 1-7 binding in the inactive-state structure compared to the Apo-state. In the active-like structure, the psi angle gradually increased from ∼70° to 90° with Ang II binding, while remaining constant at ∼60° with Ang 1-7 binding.

##### E/DRY Motif

No significant angular changes were observed in ASP141^3.49^, ARG142^3.50^, or TYR143^3.51^ across all binding states and receptor conformations.

##### CWxP Motif

For CYS268^6.47^, no significant changes occurred in the phi and psi angles with Ang II binding in the inactive-state structure. In the active-state structure, a 20° increase in the phi angle was observed after 700 ns with both ligands. The psi angle changed by ∼20° after 500 ns with Ang 1-7 binding in the inactive-state structure, while no changes occurred in the active-state structure. For TRP269^6.48^, a 10° psi angle change was observed after 500 ns with Ang 1-7 binding in the inactive-state structure, with no other notable changes. For LEU270^6.49^, increases of 10-20° in the phi angle were observed after 500 ns with Ang 1-7 binding in both receptor conformations, but no changes occurred with Ang II binding. PRO271^6.50^ showed a 30° increase in the psi angle with Ang 1-7 binding in the inactive-state structure, while other angles remained unchanged.

##### NPxxY Motif

Residues ASN314^7.49^, PRO315^7.50^, PHE316^7.51^, LEU317^7.52^, and TYR318^7.53^ showed no significant angular changes with Ang 1-7 or Ang II binding in the inactive-state structure. In the active-state structure, the phi and psi angles of these residues exhibited variations after 400-500 ns. ASN314^7.49^ and PRO315^7.50^ showed ∼20° and 10° changes, respectively, with Ang 1-7 and Ang II binding. PHE316^7.51^ exhibited 20° and 10° changes in the phi angle with Ang 1-7 and Ang II binding, respectively, while the psi angle varied only with Ang 1-7. LEU317^7.52^ and TYR318^7.53^ showed time-dependent changes in both phi and psi angles, with Ang 1-7 generally inducing larger angular shifts than Ang II.

## 6 DISCUSSION

We have demonstrated a comprehensive analysis of the ligand-specific effects of Ang II and Ang 1-7 on the activation of the AGTR2. Through long-timescale MD simulations and structural analyses, we characterized the interactions and dynamics of AGTR2 in inactive and active states. Our findings highlight key residues, structural dynamics, and signaling pathways contributing to the activation mechanism and functional role of AGTR2 within the Renin-Angiotensin System (RAS).

In the analysis of hydrogen bonds contributing to ligand-receptor interactions in inactive-state AGTR2 (Fig. 1a, Table S1), it was observed that both the PRO7-terminated Ang 1-7 and the PHE8-terminated Ang II ligands consistently formed hydrogen bonds with THR125^3.33^ through their respective PRO7 residues throughout the simulation. This finding may suggest that THR125^3.33^ serves as a critical residue for ligand recognition and receptor stabilization (64). Notably, THR125^3.33^ formed hydrogen bonds with PRO7, the terminal residue of Ang 1-7, as well as PRO7, the penultimate residue of Ang II, despite PHE8 being Ang II’s terminal residue. These interactions were preserved for the majority of the simulation, highlighting their potential importance.

A similar pattern was observed for LYS215^5.42^, which formed hydrogen bonds with the terminal residues of both ligands, PRO7 for Ang 1-7 and PHE8 for Ang II (Fig. 1a, Tables S1, S2). These bonds were also maintained for most of the simulation, suggesting that LYS215^5.42^, in conjunction with THR125^3.33^, may serve as a key residue contributing to the stabilization of inactive-state AGTR2. The presence of a conserved hydrogen bond between LYS215^5.42^ and PHE8 in both inactive and active states of AGTR2 (Table S2) further highlights its potential role in mediating ligand-receptor interactions, particularly for Ang II binding.

In addition to these key residues, ASP279^6.58^ and ASP297^7.32^ formed stable hydrogen bonds with both ligands and mediated the formation of salt bridges (Fig. 1b). These salt bridges were preserved for most of the simulation, suggesting that these residues may contribute to ligand-receptor interactions by enhancing receptor stability.

Besides these interactions, MET128^3.36^, beyond previously reported as a key residue in AGTR2 activation through its interaction with PHE8 of Ang II (33), we identified TRP100^2.60^ as a novel contributor to ligand-induced receptor stabilization through hydrophobic interactions (Fig. 1c). TRP100^2.60^ mediated a complementary role alongside MET128^3.36^ in mediating hydrophobic interactions with Ang 1-7 and Ang II in both inactive and active states, highlighting the complementary contributions of hydrophobic residues to ligand binding and receptor stability (Fig. S5).

RMSD and RMSF analyses (Fig. 2) highlighted the distinct effects of ligand binding on receptor stability and flexibility for the structural dynamics of AGTR2. Ang II binding consistently stabilized TM6 and Helix8 regions in both inactive and active states, as demonstrated by lower RMSD values compared to the Apo-form. This stabilization aligns with Ang II’s role in maintaining a structurally stable activation-ready conformation. In contrast, Ang 1-7 binding resulted in slightly higher RMSD values, indicative of enhanced receptor flexibility. RMSF data further revealed that while Ang II reduced fluctuations in TM6 and Helix8, Ang 1-7 increased motion in ICL1 and ICL2 as well as in ECL2 and ECL3, reflecting its role in facilitating structural adaptability.

Supplementary movies and Figure S5 provided direct evidence of the stable binding of both Ang II and Ang 1-7 to AGTR2 throughout the 1000 ns simulations. These findings demonstrated that both ligands remained tightly bound to the receptor in both inactive and active states. Notably, Ang II promoted minimal receptor movement, further reinforcing its stabilizing role, while Ang 1-7 exhibited more dynamic behavior, allowing for receptor flexibility.

TM-specific RMSD analysis (Fig. S4) corroborated these observations, showing that Ang II reduced fluctuations in TM6 and TM7, regions critical for signal transduction. Ang 1-7 binding, on the other hand, resulted in slightly higher RMSD in these regions, emphasizing its role as a modulator that promotes structural transitions without compromising ligand-receptor stability.

CP and DCC analyses (Fig. 3a, 3b) provided additional evidence of ligand-specific modulation of correlated receptor motions. Ang II induced synchronized positive correlations between TM3, TM6, and Helix8, indicative of stabilized inter-helical interactions that facilitate signal transduction. This observation aligns with the stabilizing effects observed in RMSD and RMSF analyses. Conversely, Ang 1-7 binding exhibited broader correlation patterns, including weaker correlations between TM6 and Helix8 and negative correlations in ICL regions, supporting its role in promoting receptor flexibility and enabling diverse signaling pathways (65). These findings were further corroborated by PCA findings (Fig. 4), which showed that Ang II may contribute to providing some of the conformational rearrangements required for signaling by stabilizing AGTR2 in both inactive and active states. This suggests that Ang II can be a candidate to promote common mechanisms of class A GPCR activation such as TM6 outward movement and stabilization of Helix8 (66,67). Ang 1-7, however, facilitated a broader conformational landscape, supporting its modulatory role in promoting receptor adaptability. Together, these findings suggest that Ang II and Ang 1-7 may have complementary roles in influencing receptor stability and flexibility, potentially aligning with mechanisms observed in class A GPCR activation.

The activation mechanism of AGTR2 was evaluated by integrating results from DRIN analyses, intracellular TM-region center of mass (COM) distance analysis, and ligand-specific effects on micro-switch motifs (68). These findings provide information about how Ang II and Ang 1-7 differentially contribute to receptor activation and align with the common activation mechanism of class A GPCRs (69).

DRIN analysis (Fig. 5, Table S3 and S4) highlighted key residues involved in signal propagation within the receptor. Ang II binding strengthened communication pathways between TM3, TM5, TM6, TM7 and Helix8, particularly through residues such as ILE132^3.40^, ASP141^3.49^, ARG142^3.50^, PRO223^5.50^, PHE265^6.44^, CYS268^6.47^, ASN314^7.49^, and TYR318^7.53^ (Fig. S6). These residues are defined as key determinants according to the common GPCR activation model, facilitating structural transitions necessary for signal transduction (69,70). In contrast, Ang 1-7 binding induced a more diffuse network of interactions, suggesting a modulatory effect that primes the receptor for alternative signaling pathways. Notably, residues such as ILE132^3.40^, ARG142^3.50^, PHE265^6.44^, CYS268^6.47^ and PRO315^7.50^ showed increased betweenness centrality with Ang 1-7, underscoring their potential role in maintaining receptor flexibility and modulating ligand-induced signaling.

The analysis of intracellular TM distances (Fig. 6) provided further evidence of ligand-specific effects on AGTR2 activation. Ang II binding in the active-state stabilized the outward displacement of TM6, a hallmark of receptor activation, while maintaining consistent TM3-TM6 distances. This stabilization may enable AGTR2 to retain an activation-ready conformation, facilitating efficient signal transduction. In the inactive state, Ang II had a more subtle impact, suggesting that its primary role lies in stabilizing the active conformation. In contrast, Ang 1-7 exhibited greater variability in TM3-TM6 and TM6-TM7 distances, reflecting its role in maintaining receptor flexibility. These observations suggest that Ang 1-7 supports the receptor’s conformational transitions, allowing it to adapt dynamically to signaling requirements without promoting full activation directly.

Dihedral angle analysis for micro-switch motifs (Fig. S7–S10) revealed ligand-induced angular changes in key motifs such as CWxP, PIF, E/DRY and NPxxY. Ang II binding promoted shifts in these motifs consistent with activation, including the outward movement of PHE265^6.44^ in the PIF motif and structural stabilization of TYR318^7.53^ in NPxxY. In contrast, Ang 1-7 induced less pronounced changes in these motifs, reflecting its modulatory role in maintaining receptor behavior rather than driving activation directly.

## 7 CONCLUSION

This study provides a detailed analysis of the ligand-specific effects of Ang II and Ang 1-7 on AGTR2 dynamics, interactions, and activation mechanism. Both ligands were shown to stably bind to inactive and active-state AGTR2 structures throughout 1-microsecond repetitive long-timescale MD simulations, highlighting their ability to influence receptor structural dynamics. While Ang II and Ang 1-7 contribute to the stabilization of active-state AGTR2 conformations, Ang II appears to have a more pronounced effect, potentially prolonging the receptor’s active state and enhancing the likelihood of intracellular G-protein or β-arrestin binding.

However, our findings on the transitions between inactive and active states remain inconclusive, as the simulations did not fully capture these transitions. This limitation underscores the need for longer MD simulations extending beyond 1 μs, as well as experimental studies, particularly for AGTR2, which currently lacks crystal structures of its inactive or intermediate states. Despite these constraints, the results of this study provide valuable insights into the activation mechanism of AGTR2 and offer a framework for further research into its unique signaling roles within the RAS. These findings also present potential avenues for therapeutic targeting of GPCR-mediated cardiovascular regulation.

## DATA AND CODE AVAILABILITY

The codes used for analyzing micro-switch dihedral angles and hydrophobic interactions during molecular dynamics simulations are publicly available on GitHub at the following links: https://github.com/eygpcr/microswitches_dihedral_angles https://github.com/eygpcr/hydrophobic_interactions

Trajectory files of the simulations and related data for AGTR2 can be accessed through the GPCRmd database using the following identifiers: PDB code: 6JOD, Uniprot ID: P50052 https://www.gpcrmd.org/dynadb/search/

## SUPPORTING MATERIAL

Supporting Materials and Methods, which illustrate detailed analyses, including ligand-receptor interactions, structural dynamics and activation mechanisms, include ten figures, four tables and a data file for the movies (https://zenodo.org/records/14697886).

## AUTHOR CONTRIBUTIONS

E.Y. designed and performed the research and wrote the manuscript. S.D., E.E., and N.Y. designed the research and wrote the manuscript.

## ACKNOWLEDGMENTS

This work was supported by Akdeniz University Scientific Research Projects Found Grant No. TDK-5449, 2021.

