## Supplemental Materials for "Ligand Binding Effects on Activation of Type-2 Angiotensin II Receptor"

**SUPPORTING MATERIAL**

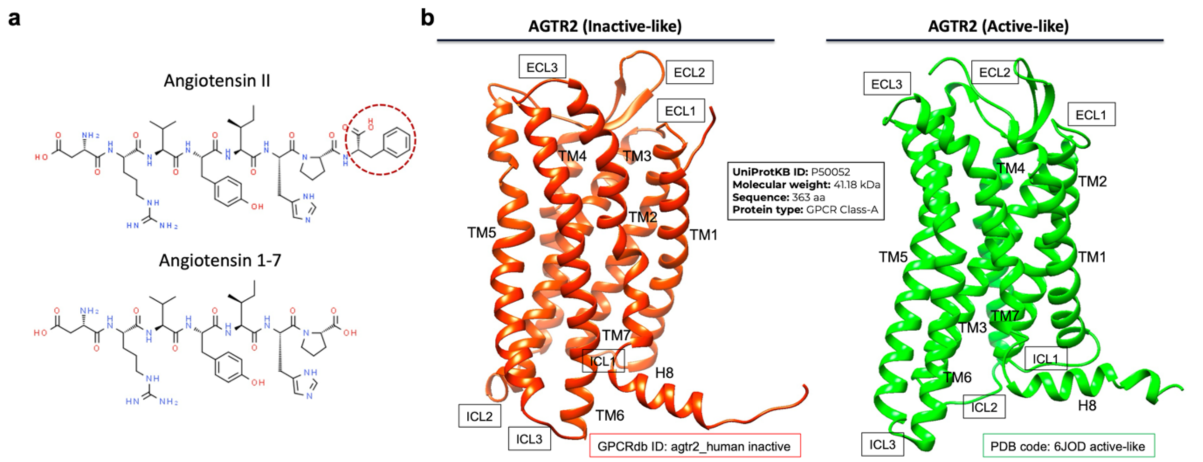


Fig S1. a) Chemical structure of Ang II and Ang 1-7 peptides, b) Inactive-like and active-like structures of AGTR2.

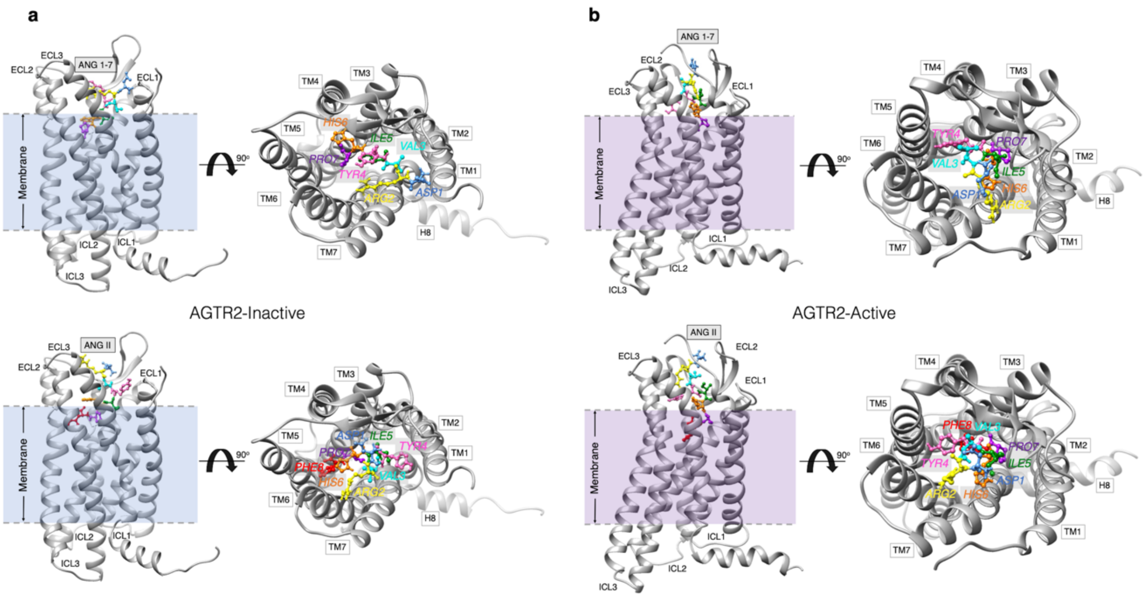


Fig S2. Interaction poses of AGTR2 with ligands according to best docking scores. Binding interactions of Ang II and Ang 1-7 ligands with a) inactive and b) active structure of AGTR2.

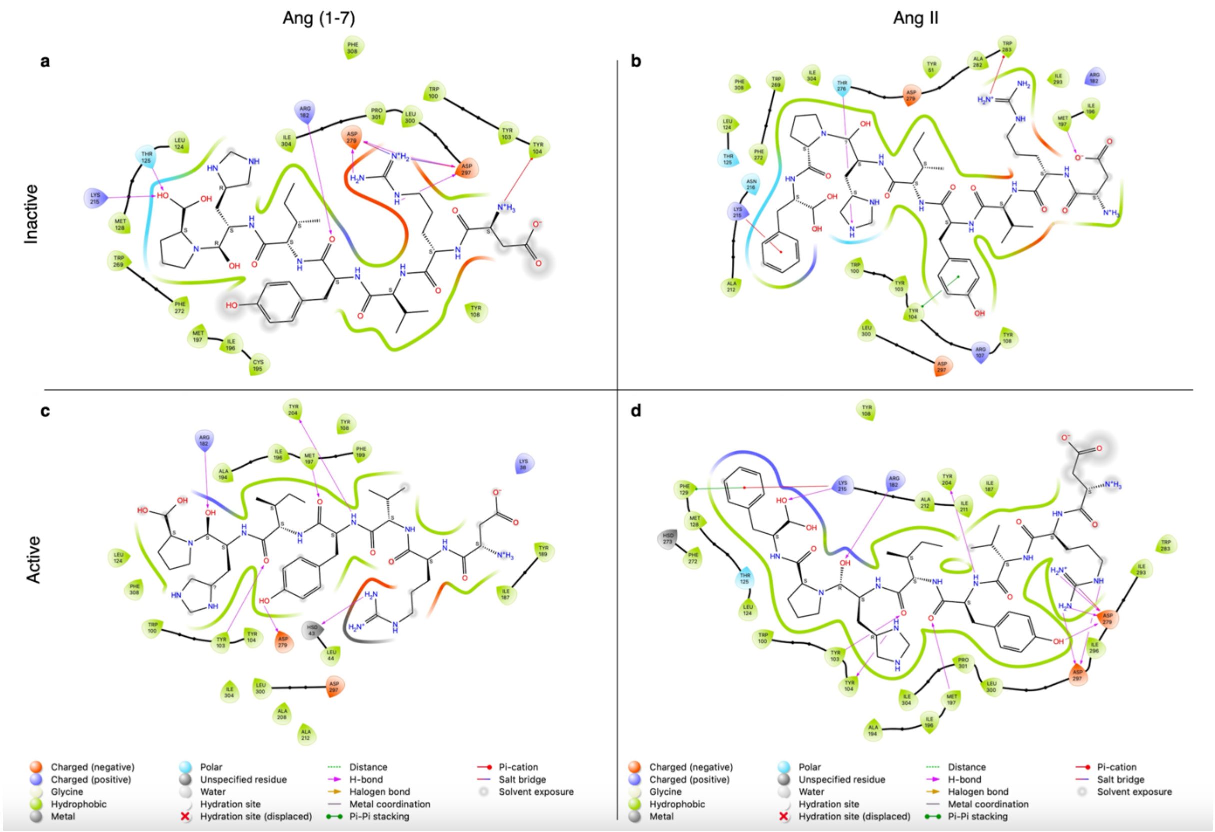


Fig S3. 2D ligand interactions of AGTR2 with ligands according to best docking scores.

Table S1. Hydrogen bond interactions of the inactive-state AGTR2 with Ang 1-7 and Ang II ligands throughout the simulation. (The red color represents the residues of the ligands.)

| Donor | Acceptor | Frames |
|---|---|---|
| THR125-Side | PRO7-Side | 1293 |
| ARG2-Side | ASP297-Side | 888 |
| ARG2-Side | ASP279-Side | 743 |
| LYS215-Side | PRO7-Side | 657 |
| SER40-Main | ASP1-Side | 441 |
| LYS42-Side | ASP1-Side | 369 |
| ARG182-Side | TYR4-Main | 358 |
| ASP1-Main | ASP41-Main | 212 |
| ASP1-Main | ASP41-Side | 209 |
| TYR4-Side | ASP279-Side | 172 |
| HIS6-Side | THR276-Side | 166 |
| TYR108-Side | ASP1-Main | 109 |
| TYR104-Side | ASP1-Main | 77 |
| ASP1-Main | ASP297-Side | 50 |

| Donor | Acceptor | Frames |
|---|---|---|
| LYS215-Side | PHE8-Side | 945 |
| THR125-Side | PRO7-Main | 811 |
| ARG182-Side | HIS6-Main | 803 |
| ASP1-Main | ASP297-Side | 759 |
| ARG2-Side | ASP279-Side | 486 |
| TYR103-Side | TYR4-Main | 344 |
| ASP1-Main | ASP279-Side | 107 |
| TYR108-Main | TYR4-Side | 52 |

Table S2. Hydrogen bond interactions of the active-state AGTR2 with Ang 1-7 and Ang II ligands throughout the simulation. (The red color represents the residues of the ligands.)

| Donor | Acceptor | Frames |
|---|---|---|
| ASP1-Main | GLU188-Side | 449 |
| ARG185-Side | ASP1-Side | 172 |
| ARG182-Side | PRO7-Side | 126 |
| LYS38-Side | ASP1-Side | 119 |
| LYS215-Side | PRO7-Side | 62 |
| GLU188-Main | ASP1-Main | 60 |
| ASP1-Main | ASP297-Side | 55 |

| Donor | Acceptor | Frames |
|---|---|---|
| TYR4-Main | TYR204-Side | 867 |
| LYS215-Side | PHE8-Side | 750 |
| CYS35-Main | ASP1-Side | 714 |
| ARG2-Side | ASP279-Side | 631 |
| ARG2-Side | ASP297-Side | 465 |
| ARG182-Side | HIS6-Main | 456 |
| MET197-Main | TYR4-Main | 338 |
| ARG182-Side | PHE8-Side | 146 |
| ASP1-Main | GLU188-Side | 144 |
| ASP1-Main | GLN37-Side | 116 |
| ARG182-Side | ILE5-Main | 111 |
| SER36-Side | ASP1-Side | 69 |
| GLN37-Side | ASP1-Side | 66 |
| LYS215-Side | TYR4-Side | 56 |

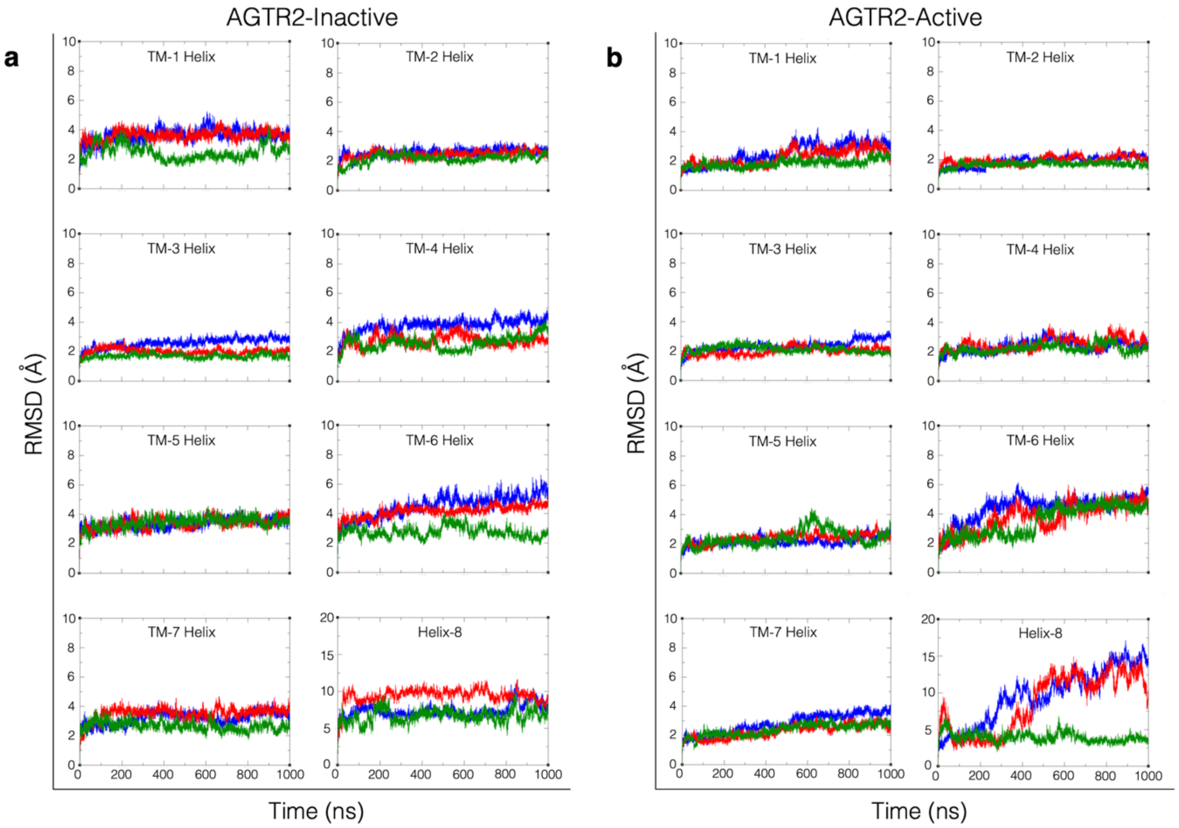


Fig S4. RMSD analyses of the TM helical structures of AGTR2. a) Inactive-state and b) active-state RMSD change plots of TM helices of Apo-AGTR2, Ang (1-7)-AGTR2 and Ang II-AGTR2 complex structures. (Blue color represents Apo-AGTR2, red color Ang (1-7)-AGTR2 and green color Ang II-AGTR2 complex structures.)

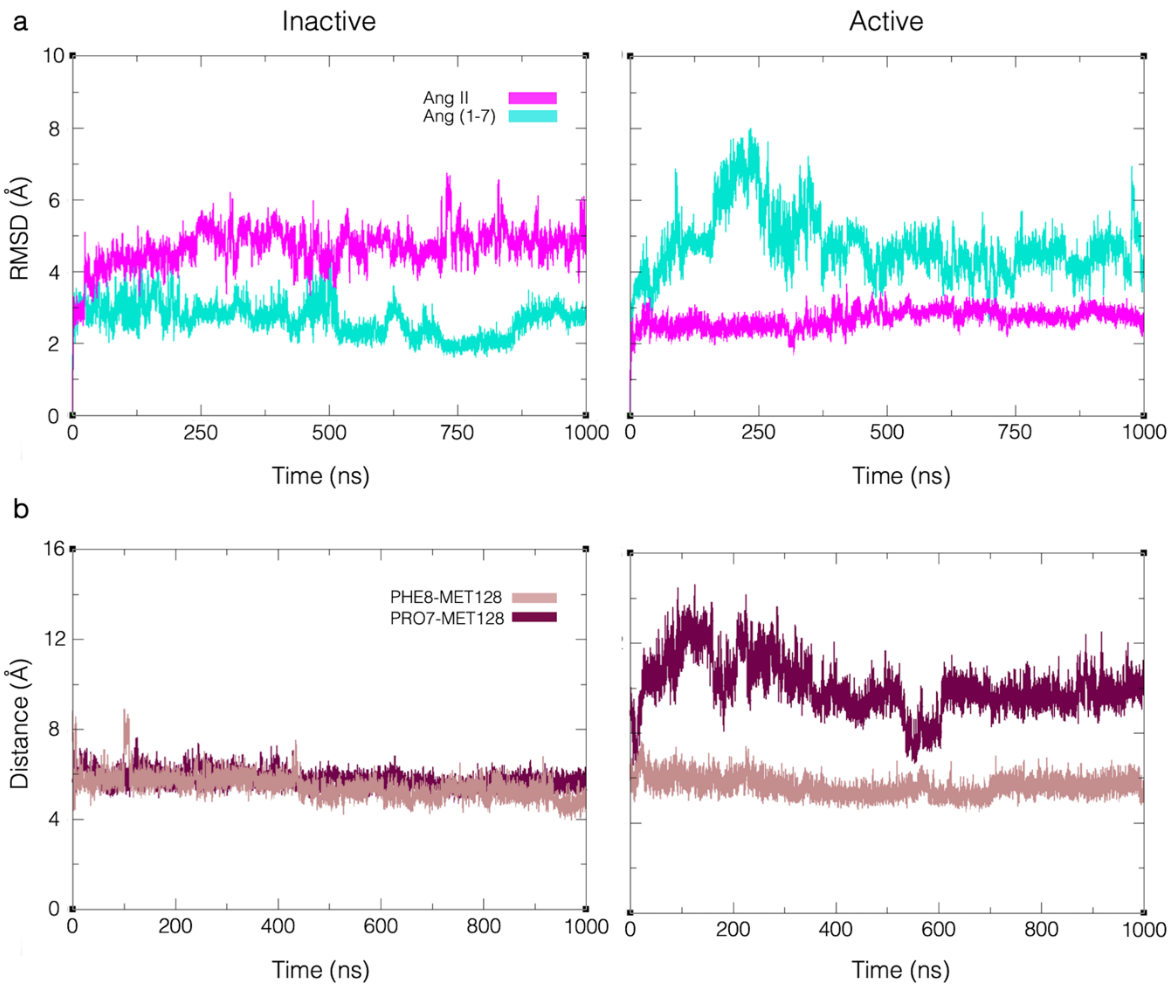


Fig S5. RMSD and distance analyses of the Ang II and Ang 1-7 ligands. a) RMSD analyses of Ang II and Ang 1-7 ligands both inactive and active states during simulation. b) Distance analyses of between last residues of ligands and MET128$^{3.36}$ key residue responsible for activation.

Table S3. List of AGTR2 residues for both increasing and decreasing BC and mean L in the Ang 1-7 binding state. (Bold expressions represent micro-switches residues.)

| | Standard deviation | Residues with significant increase in BC | Residues with significant decrease in BC | Residues with significant increase in average L | Residues with significant decrease in average L |
|---|---|---|---|---|---|
| Inactive | **±3$\sigma$** | SER79, **ARG142**, ILE249, THR250, GLN253, VAL254, MET257, ALA260, VAL261, ALA264, **PHE265**, LYS328, VAL332, LEU340 | VAL262, **CYS319** | LEU151, SER152, PRO336, ILE337, THR338 | SER40, ASP41, LYS42, HIS43, PRO75, TYR108, LYS203, TYR204, ILE249, THR250, ASP252, GLN253, VAL254, MET257, SER331, VAL332, PHE333, TRP339, LEU340, GLN341 |
| | **±2$\sigma$** | ILE83, LEU86, LEU94, TYR103, CYS117, GLY121, LEU124, THR125, ALA130, **ILE132**, PHE134, TYR148, PHE199, GLY219, ALA258, **CYS268**, LEU275, THR276, PHE277, PHE325 | ALA259, PHE320, GLY322, ASN323, GLN326, ARG330, ILE337 | GLN72, LYS73, PHE150, GLU202, THR241 | LEU44, GLN159, PHE199, ARG251, LEU255, LYS256, ALA258, ALA260 |
| Active | **±3$\sigma$** | LEU124, GLN153, ILE304, PHE320, PHE325 | ILE47, ALA87, LEU94, THR96, LEU97, TYR103, TYR104, PHE123, LEU126, ALA130, ILE135, SER139, ALA212, ASN216, VAL262, **TYR318**, **CYS319**, LEU329 | GLY74, PRO75, LEU151, GLN153, ARG154 | PRO39, ASP41, LYS42, TYR108, PHE112, ILE187, GLU188, TYR189, LEU190, ALA194, ILE196, THR241, ASN242 |
| | **±2$\sigma$** | GLY121, **CYS268**, LEU300, **PRO315** | ILE54, PHE69, GLY74, ALA89, TRP110, PHE120, PHE129, PHE134, **ARG142**, VAL167, CYS195, ALA198, PHE199, GLY219, LYS256, **PHE265**, HIS273, ASN310 | LYS246, ASN323, LEU329, ARG330, PHE333 | CYS35, SER36, SER40, HIS43, LEU44, ARG107, ASP109, LEU111, PRO114, MET116, GLY121, LEU124, SER152, ASP183, VAL184, ARG185, THR186, GLY191, VAL192, ASN193, MET197, GLU202, ARG235, LEU239, ASP297 |

Table S4. List of AGTR2 residues for both increasing and decreasing BC and mean L in the Ang II binding state. (Bold expressions represent micro-switches residues.)

| | Standard deviation | Residues with significant increase in BC | Residues with significant decrease in BC | Residues with significant increase in average L | Residues with significant decrease in average L |
|---|---|---|---|---|---|
| Inactive | $\pm 3\sigma$ | SER79, ILE83, LEU86, THR125, **ILE132**, CYS137, **ARG142**, TYR189, PHE199, VAL254, MET257, ALA264, **PHE265**, THR276, LEU300, ILE304, LEU305, THR309, VAL313, **TYR318**, CYS319, VAL321, GLY322, PHE325, GLN326, LEU329, ARG334 | NONE | NONE | SER40, ASP41, LYS42, HIS43, LEU44, ASP45, ALA46, ILE47, GLN72, ARG107, TYR108, ASP109, PRO149, PHE150, GLN153, ARG185, THR186, ILE187, GLU188, TYR189, LEU190, GLY191, VAL192, ALA194, PHE199, PRO201, GLU202, LYS203, TYR204, ALA205, LEU239, LYS240, THR241, ASN242, SER243, TYR244, GLY245, LYS246, ASN247, ARG248, ILE249, THR250, ARG251, ASN288, SER289, CYS290, GLU291, ALA294, GLN327, ARG330, SER331, VAL332, PHE333, ARG334, VAL335 |
| | $\pm 2\sigma$ | ILE47, LEU50, TYR82, **ASP90**, TRP100, PHE133, MET138, SER139, SER145, SER175, ARG182, SER208, GLY219, **PRO223**, ILE227, ARG251, ALA258, VAL261, **CYS268**, GLY307, ASN310, **ASN314**, **PHE316**, LYS328, VAL332 | SER174, PRO177 | NONE | PRO48, ILE49, LEU50, TYR52, ILE53, VAL64, TYR67, LEU68, PHE69, CYS70, CYS71, GLY74, PRO75, LYS76, TYR103, TYR104, SER105, TYR106, TRP110, LEU111, PHE112, GLY113, PRO114, VAL115, GLN144, SER145, VAL146, ILE147, TYR148, LEU151, SER152, ARG154, ASN156, TRP158, GLN159, TYR162, ASP183, VAL184, ASN193, CYS195, ILE196, MET197, ALA198, PRO200, GLN206, TRP207, SER208, ALA209, PHE232, GLY233, ILE234, ARG235, LYS236, HIS237, LEU238, ASP252, GLN253, VAL254, LEU255, LYS256, MET257, GLY285, VAL286, ILE287, VAL292, ILE293, VAL295, ILE296, ASP297, LEU298, ALA299, **CYS319**, ASN323, ARG324, GLN326, LYS328, LEU329 |
| Active | $\pm 3\sigma$ | PRO39, SER40, LEU68, VAL78, TYR82, GLY121, SER122, LEU124, **MET128**, SER131, SER208, LYS215, LEU300, ILE304, **ASN314**, VAL332 | **CYS319**, PHE320, LEU329, ARG330, LEU340 | GLY74, PRO75, PHE150, LEU151, ARG154, SER243, TYR244, GLY245, LYS246, ASN247, ARG248, ILE249, THR250, ARG251, GLN253, VAL254 | LYS42, VAL332, VAL335, PRO336 |
| | $\pm 2\sigma$ | PHE59, ILE63, VAL64, VAL65, THR67, ILE81, ASN85, LEU86, LEU93, PHE134, **ASP141**, ARG155, SER175, ILE196, LEU255, GLY307, PHE325, VAL335 | ILE47, THR96, TYR104, TRP110, PHE120, GLN144, ALA198, PHE199, GLN326 | VAL66, PHE69, CYS70, GLN72, LYS73, GLY113, PRO114, VAL115, VAL119, TYR143, VAL146, ILE147, PRO149, SER152, GLN153, ASN156, GLN159, TYR162, ILE163, LYS236, HIS237, LEU238, LYS240, ASP252, ASN288, **CYS319**, ARG330 | PRO39, SER40, LEU111, PHE112, LEU124, TYR189, ILE196, SER331 |

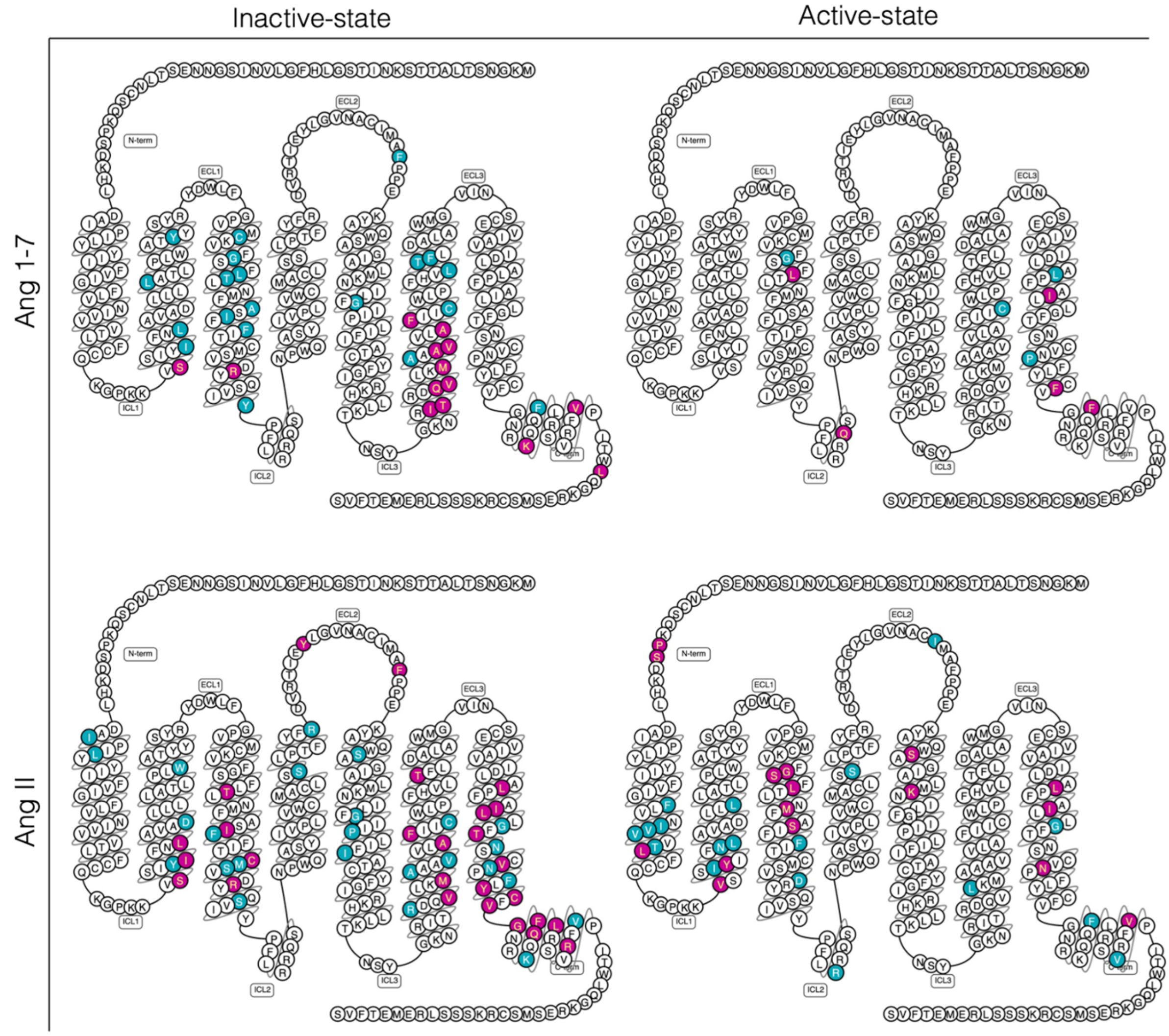


Fig S6. Snake diagrams of AGTR2 residues in inactive and active states showing increasing BC values for both Ang 1-7 and Ang II binding states. (Magenta color represents increasing BC values for residues exceeding 3σ, while cyan color represents increasing BC values for residues exceeding 2σ.)

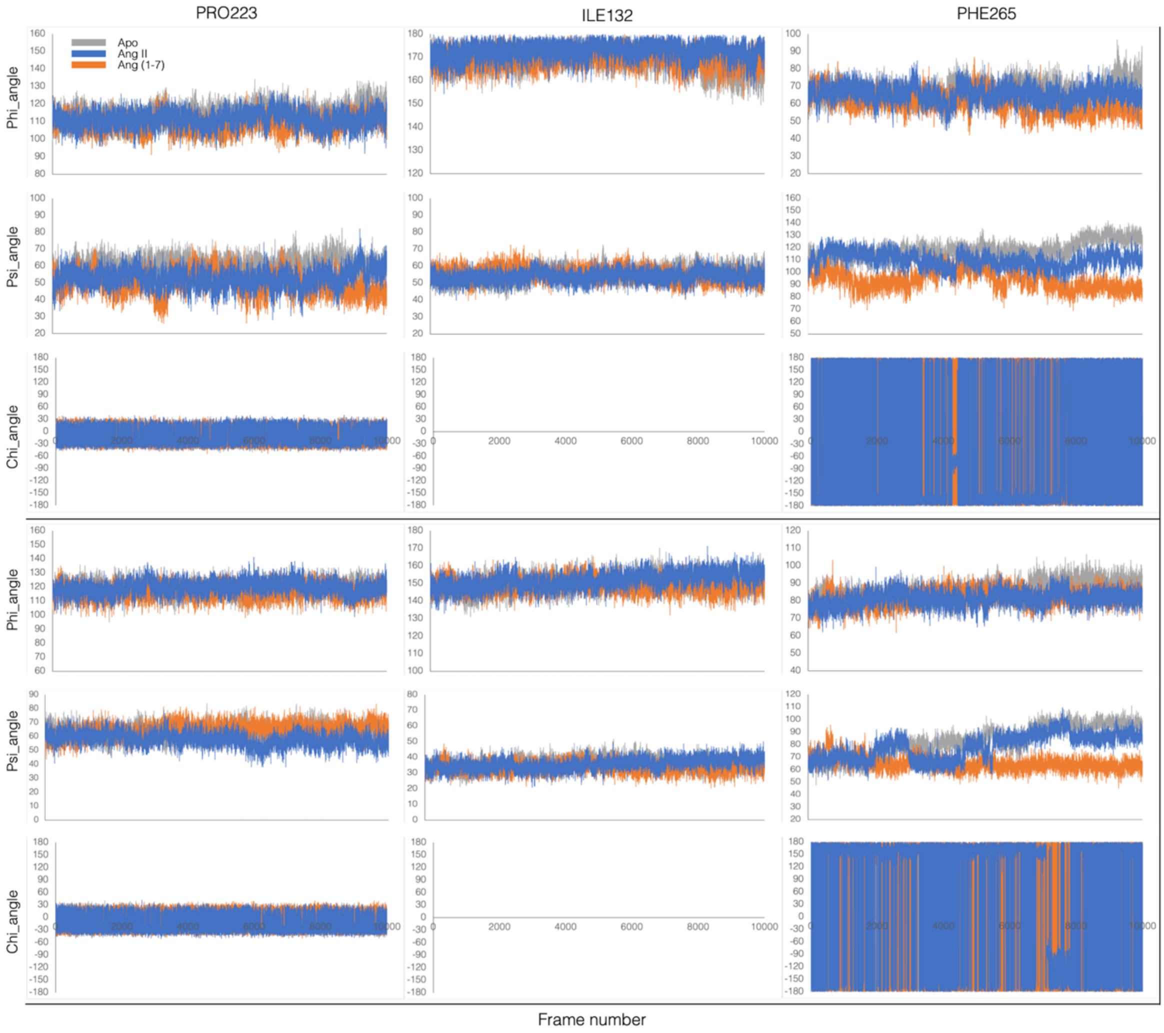


Fig S7. Dihedral angle changes (phi, psi, and chi) of the PIF motif (PRO223$^{5.50}$-ILE132$^{3.40}$-PHE265$^{6.44}$) during simulations in the ligand-bound and unbound states of inactive and active-like AGTR2.

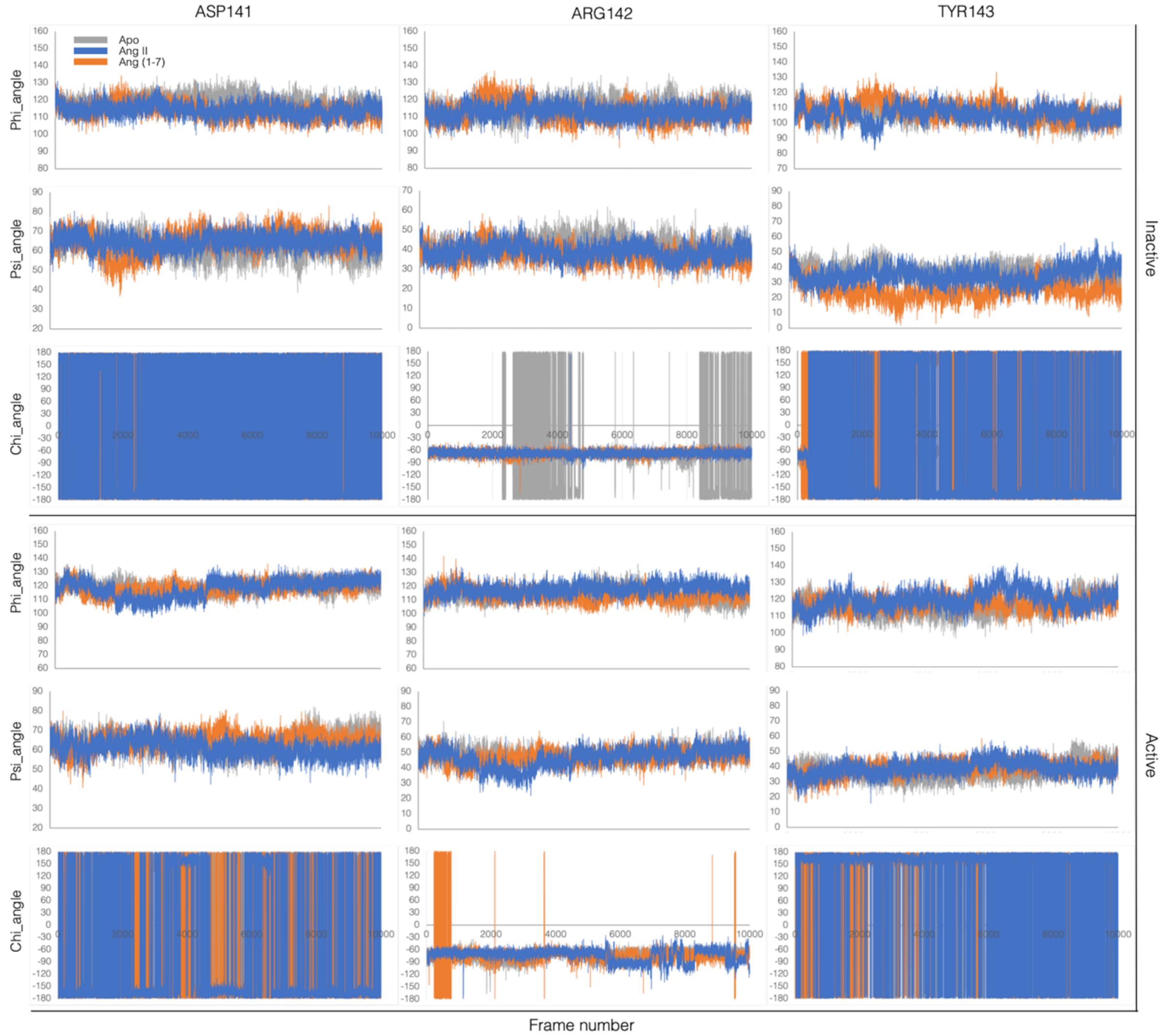


Fig S8. Dihedral angle changes (phi, psi, and chi) of the E/DRY motif (ASP141$^{3.49}$-ARG142$^{3.50}$-TYR143$^{3.51}$) during simulations in the ligand-bound and unbound states of inactive and active-like AGTR2.

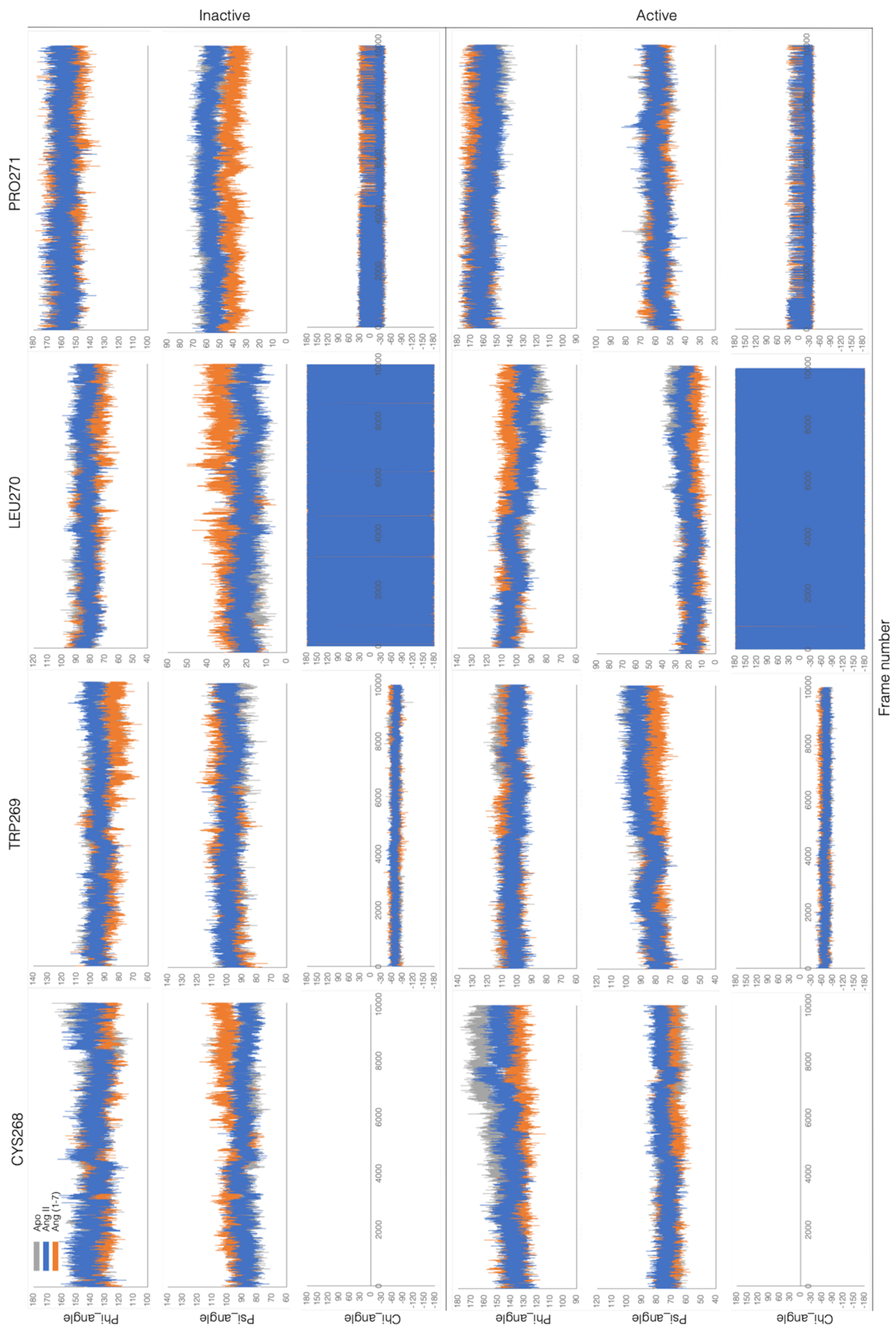


Fig S9. Dihedral angle changes (phi, psi, and chi) of the CWxP motif (CYS268$^{6.47}$-TRP269$^{6.48}$-LEU270$^{6.49}$-PRO271$^{6.50}$) during simulations in the ligand-bound and unbound states of inactive and active-like AGTR2.

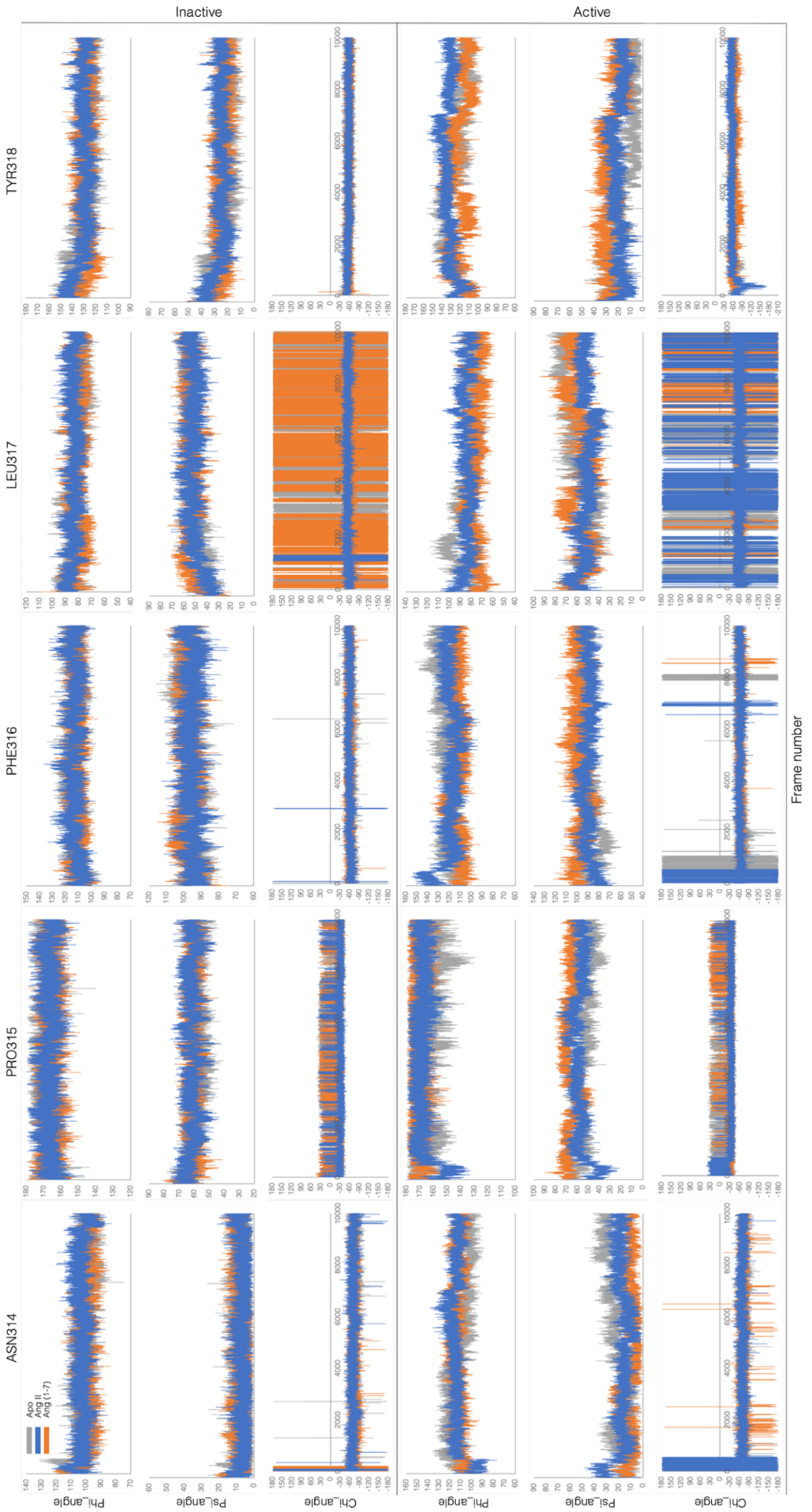


Fig S10. Dihedral angle changes (phi, psi, and chi) of the NPxxY motif (ASN314$^{7.49}$-PRO315$^{7.50}$-PHE316$^{7.51}$-LEU317$^{7.52}$-TYR318$^{7.53}$) during simulations in the ligand-bound and unbound states of inactive and active-like AGTR2.

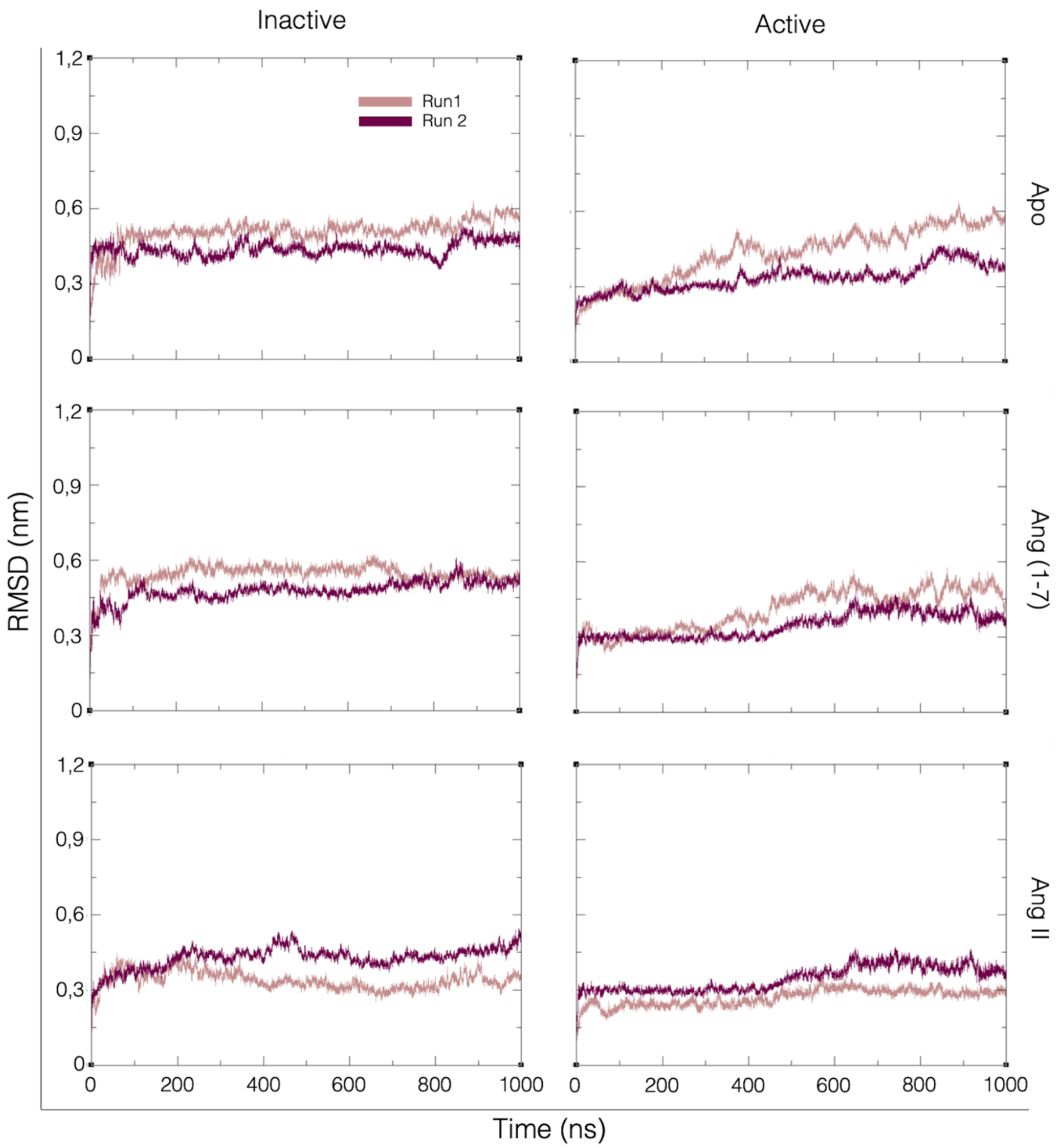


Fig S11. RMSD analyses of two different replicates of AGTR2 in both inactive and active states with and without ligand binding.
